# Effect of the environment, predation and life history on reproductive synchrony in Perissodactyla

**DOI:** 10.64898/2026.06.09.731243

**Authors:** Lucie Thel, Elise Huchard, Dieter Lukas, Bernard Godelle, Jules Dezeure, Jan A. Venter

## Abstract

Synchronising birth timing with the adequate environmental conditions as well as with congeners can affect juvenile survival and at a later stage, recruitment. In the case of endangered species, understanding the factors driving birth phenology is therefore critical to implement effective conservation and reintroduction policies. Currently, 75% of the Perissodactyla species (odd-toed ungulates) are threatened and some even already had to be reintroduced in their natural habitat. Using a comparative analysis framework, we investigated the effect of nine environmental, life history and predation factors on the synchrony of births in 27 populations of wild Perissodactyla across Africa and Asia. We confirmed the positive effect of environmental seasonality and latitude, as well as the negative effect of environmental productivity on birth synchrony, but we found no significant effect of environmental unpredictability. Life history traits had limited effects on birth synchrony in this order: the intensity of daily reproductive effort and pace of life were non-significant, and migratory behaviour had an unexpected positive association with birth synchrony, suggesting that the link between migration and reproduction might be more complex than the relief of environmental constraints initially expected. We found a positive effect of group size on birth synchrony, in line with the swamping hypothesis to reduce predation risk on neonates, but we found no strong support for a positive effect of predation exposure, measured as a qualitative index combining species-specific sensitivity to predation and site-specific predator abundance. Overall, our findings suggest that individuals adjust reproductive cycles in response to local ecological and social conditions, suggesting that conservation efforts such as reintroductions could benefit from taking into account the extent of the differences between the site of origin and the reintroduction site.

## Introduction

Timing reproduction adequately is critical for offspring survival (Feder et al. 2008, Lee et al. 2017), and consequently affects population dynamics and maintenance (Gaillard et al. 2000, Raithel et al. 2007). Synchronising the reproductive cycle with the period of food abundance allows individuals to optimise energetic processes, particularly when resources are not homogeneously spread out during the year, as is the case in most environments (Bronson 1989). However, numerous additional factors are involved in determining the date of birth besides energetic constraints, whether intrinsic such as life history traits, or extrinsic such as predation exposure.

Food resources act as the predominant factor driving reproductive timing (e.g. Sinclair et al. 2000 in ungulates, Di Bitetti and Janson 2000 in primates, Heideman and Utzurrum 2003 in nectarivorous bats). In seasonal, predictable environments, reproductive timing should ensure that the most costly part of the reproductive cycle (late gestation and early lactation in large herbivores; Sadleir 1982) matches the peak of food availability and/or quality (Bronson 1989), thus favouring reproductive seasonality (*sensu* Burtschell et al. 2026). Climatic conditions such as temperature and humidity simultaneously act on the reproductive timing by affecting neonate survival, which is exposed to energy loss because of its limited ability to thermoregulate (Grovenburg et al. 2012, Mattisson et al. 2022). In unpredictable environments (Colwell 1974), to the contrary, reproductive seasonality could lead to missed breeding opportunities or even unsuccessful breeding in the case of a mismatch between adequate environmental conditions and the reproductive cycle (Burtschell et al. 2026). Finally, in highly productive systems where energy acquisition and conservation are not a constraint, reproduction should be less seasonally constrained and rather driven by other factors.

Life history traits also contribute to shaping the level of reproductive seasonality. Species with slow pace of life requiring long gestation and lactation periods for foetal and juvenile development (Kihlström 1972) are more likely to spread the reproductive costs throughout the year and breed non-seasonally (Ahrestani et al. 2012). Species capable of behavioural adjustments might also be better able to shift the timing of their births to avoid costs or to gain from exploiting favourable conditions (Burtschell et al. 2026). For instance, migratory behaviour (e.g. Aikens et al. 2021) should allow individuals to reduce their dependency to temporally constrained food availability and relax the need for a seasonal reproduction, or could extend the period during which resources are abundant, thereby favouring a seasonal reproductive strategy to take advantage of the most favourable period of the year.

Although energetic considerations are key drivers of reproductive seasonality, certain species display pronounced synchrony at low latitudes (e.g. blue wildebeests (*Connochaetes taurinus*) in the Ngorongoro, Tanzania; Estes 1976), while others exhibit various levels of reproductive synchrony despite living in the same environment (e.g. large herbivores guild in the Serengeti, Tanzania; Sinclair et al. 2000). These observations highlight the existence of additional factors influencing reproductive synchrony (Ims 1990a). Namely, most populations of large herbivores interact with a guild of predators, a main cause of neonate mortality (Linnell et al. 1995, Severud et al. 2019). Ims’ (1990b) theoretical work suggests that specialist predators (which target exclusively certain prey species) should favour reproductive synchrony among their prey via the swamping strategy (Darling 1938): a high concentration of prey appearing at the same time limits individual vulnerability to predation since predators are maintained at a low density due to seasonal resources and can only catch and handle a limited number of prey at a time. Similarly, prey species living in large aggregations should display synchronous parturitions to benefit from the swamping effect (Sinclair et al. 2000). However, a non-synchronous reproduction could be more advantageous when confronted to generalist predators, which change prey according to their temporal availability and can thus maintain high densities (Ims 1990b). Many simultaneous births among their prey would lead predators to switch their diet from alternative prey species during the rest of the year toward the newborns at that specific period, dramatically increasing predation risk on these neonates. Deriving from these evolutionary hypotheses, prey populations exposed to predators to which they are highly vulnerable should also flexibly adjust parturition to give birth more synchronously and similarly, benefit from the swamping effect.

Perissodactyla, which comprise 16 species in the wild, are exposed to varied environmental and predation conditions in their natural habitats. They inhabit African and Asian plains (mostly *Equidae* and African *Rhinocerotidae*) and South American and Asian tropical and equatorial forests (mostly *Tapiridae* and Asian *Rhinocerotidae*). They display a variety of birth seasonality, from a few weeks in summer for the Asian *Equidae* such as the kiang (*Equus kiang*; Schaller 1998) and the onager (*Equus hemionus*; Shah and Qureshi 2007), to year-round parturitions for some populations of plains zebra (*Equus quagga*; Thel et al. 2025) and black rhinoceros (*Diceros bicornis*, Hrabar and du Toit 2005) in Africa.

The long life-span and slow generation time of Perissodactyla reduce their ability to respond to rapid changes, making them particularly vulnerable to threats such as climate change (Hoffmann and Sgrò 2011). Additionally, multiple species such as the African wild ass (*Equus africanus*) and all species of *Rhinocerotidae* are under severe poaching pressure (Kebede et al. 2014, Clements et al. 2020), while all species of *Tapiridae* are threatened by deforestation and habitat fragmentation (e.g. Magintan et al. 2012). Currently, 75% of the Perissodactyla species are threatened and four are critically endangered (IUCN Red List, https://www.iucnredlist.org/). Understanding reproductive biology plays an important role in designing efficient conservation plans, including reintroduction operations. In reintroduction sites, several years can be necessary for a population to adjust birth phenology to local conditions, potentially reducing juvenile survival (Whiting et al. 2011, Whiting et al. 2012). However, Perissodactyla have been overlooked in comparison with Artiodactyla when it comes to their reproductive biology in the wild (e.g. Ryan et al. 2007, Kourkgy et al. 2016, Lee et al. 2017, Laforge et al. 2023, Thel et al. 2026). Considering this knowledge gap and the current conservation status of this order, there is an urgent need to better understand their birth phenology and ecological drivers.

Using a comparative analysis in a Bayesian framework based on published phenology of births, we investigated the effect of nine environmental, life history traits and predation factors on birth synchrony in 10 species of Perissodactyla, spanning 27 wild populations across Africa and Asia. We expected that environmental factors such as latitude and environmental seasonality should increase birth synchrony, while environmental productivity and unpredictability should decrease birth synchrony. Additionally, we expected life history traits such as daily reproductive effort and pace of life to increase birth synchrony, while migratory behaviour should reduce birth synchrony if it buffers resources seasonality or else increase it. Finally, group size and predation exposure should increase birth synchrony.

## Methods

### Birth phenology

We identified the species of non-extinct wild Perissodactyla using the Mammal Diversity Database (https://www.mammaldiversity.org/, search date: April 2025, *n* = 16). Although the Przewalski’s horse (*Equus ferus*) has been extinct in the wild since the 1960s, we decided to include this species as it has been successfully reintroduced and monitored in its natural habitat since the 1980s (Xia et al. 2014). We then conducted a thorough species-specific search in Google Scholar (https://scholar.google.fr/, search date: October 2025) using a unique combination of two sets of key words: i) the scientific, common or less common/old species name, when adequate (e.g. the greater one-horned rhinoceros (*Rhinoceros unicornis*) is also called Indian rhinoceros), ii) one of six reproduction key words: “reproduction”, “birth”, “parturition”, “foaling” (or “calving”, when appropriate), “breeding”, or “newborn”. We scanned the 20 first results for each search. Considering the redundancy of results between the different searches for a given species, we are confident that the number of studies missed in the search is likely to be very low. Because the number of studies on Perissodactyla breeding phenology is low and the number of births per month is a first-order variable (i.e. leaves little room for misinterpretation or errors during data collection), we did not restrict the search to peer-reviewed literature and included books as well as technical, master and PhD reports written in English and published after 1900 (Supporting Information 1). We identified a total of 66 potentially relevant studies, among which 46 provided a complete or partial monthly distribution of births.

For the analyses, we only included populations monitored for at least one year, and in which at least 12 births were recorded during the monitoring period (see details on data selection in Supporting Information 1). For each study, we collected the number of births (alternatively, proportion of births and total sample size) per month. When necessary, we extracted this data from graphs using WebPlotDigitizer (Drevon et al. 2017). Following current practices (Thel et al. 2022, Dezeure et al. 2024, Burtschell et al. 2026), we computed the mean vector length *r* from circular statistics to describe birth synchrony (*n* = 27 populations). Each month was considered as an angle of 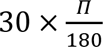 radians. When births were perfectly evenly distributed, *r* = 0, and when they were all clustered in the same month, *r* = 1.

### Phylogenetic correlation

Following the process described by Burtschell et al. (2026), we retrieved a credible set of 1000 trees of all species of Perissodactyla (https://vertlife.org/phylosubsets/, search date: November 2025). We used TreeAnnotator (BEAST, version 10.5.0; Drummond et al. 2012) to generate a maximum clade credibility (MCC) tree (median node heights and burn in of 250 trees). We trimmed the tree to match the species in our dataset. We tested the strength of the phylogenetic signal in reproductive synchrony among each dataset using Blomberg’s *K* using the “phylosig” function with *n* = 1000 randomisations (“phytools” package; Revell 2024). A *K* value close to 0 means that closely related species are not similar to each other, while a *K* value close to 1 means that they are. As we did not detect any phylogenetic signal that would indicate significant long-term evolutionary processes among the species included in the study (Blomberg’s *K* = 0.27, *p* = 0.11), we focussed on contemporary data and their ecological effect on birth synchrony, with regard to both environmental predictors and predation exposure.

### Environmental predictors

When reported in the study, we extracted the latitude of the study site. Otherwise, we either used the coordinates provided by another study in the same study site or searched the name of the study site on <u>latitude.to</u> and used the corresponding GPS coordinates. When the study encompassed multiple geographically close study sites, we used the mean GPS coordinates or the GPS coordinates of the main site (n = 2 studies, Supporting Information 1).

To extract site-specific environmental predictors, we drew a rectangle around the border of each site delineated by the extreme longitudes and latitudes of the geographical unit (e.g. private or national park, administrative region or district). We retrieved the mean bi-weekly Enhanced Vegetation Index (EVI) in this area between March 2000 and December 2024, from Google Earth Engine (https://earthengine.google.com/) using the moderate resolution imaging spectroradiometer MOD13A1.061 Terra Vegetation Indices 16-Day Global 500m (Didan and Huete 2021). The EVI was designed to improve the Normalized Difference Vegetation Index (NDVI) by reducing biases due to atmospheric and canopy background effects (Huete et al. 1997), and maintaining sensitivity over high biomass locations (Gao et al. 2000). We used linear interpolation (function “na.approx” of the package “zoo”; Zeileis and Grothendieck 2005) to estimate daily EVI values. Some study sites in Mongolia and China (*n* = 4, 1.6% of total sample size) were characterised by negative EVI values, denoting barren soil or snow cover. In both instances, negative values correspond to the absence of vegetation, and we set these values to 0 for further analyses.

We used the EVI as a proxy of food resources, as satellite-derived greenness indices are commonly used in large herbivore studies (e.g. Ryan et al. 2007, English et al. 2012, Laforge et al. 2023). For each study site, we calculated the mean monthly EVI across the 25 years of environmental data available. We then estimated environmental productivity, seasonality and unpredictability following the methods described by Dezeure et al. (2024). The EVI was decomposed into three components: *EVI_m,y_* = *C* + *S_m_* + *NS_m,y_* = *C*, where *m* is the month of the year (January to December) and *y* is the year (2000 to 2024). *C* is a constant corresponding to the mean monthly EVI over the full time series, *S_m_* is the seasonal component of the monthly EVI and *NS_m,y_* is the non-seasonal component of the monthly EVI, defined as:

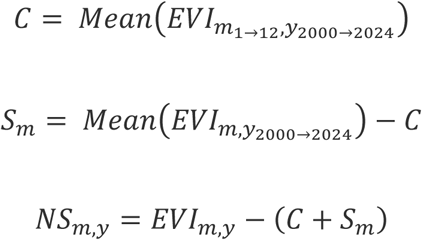

From there, the three environmental predictors were defined as:

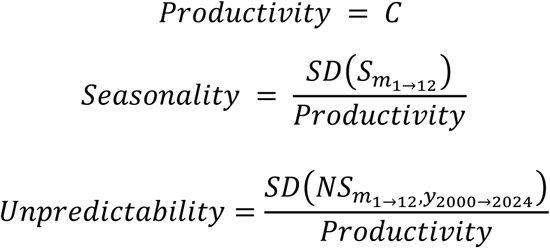

The higher the *Seasonality* index, the more seasonal the EVI fluctuations, and the higher the *Unpredictability* index, the more unpredictable the EVI fluctuations.

### Life history traits

To determine the pace of life of each species, we collected life history traits from the database of Myhrvold et al. (2015): birth mass, female adult mass, gestation length, lactation length, interbirth interval and female age at maturity. When information was missing for certain species, we conducted species-specific searches and used complementary sources (see details in Supporting Information 2). We then conducted a principal component analysis (PCA) including these life history traits while correcting for differences in body mass (Bielby et al. 2007). We used the residuals of the regression of each life history trait according to the log-transformed female adult mass in the PCA. The first axis of the PCA explained 66% of the variance, and the species with the slowest pace of life relative to its mass, the Sumatran rhinoceros (*Dicerorhinus sumatrensis*), displayed the highest value. The second axis explained 21% of the total variance (Supporting Information 2). To simplify the interpretations, we used the coordinates of the first axis multiplied by -1 as a proxy for the pace of life, so that the species with the highest value of this variable displays the fastest pace of life (relative to its mass) and the species with the lowest value displays the slowest pace of life.

To quantify the intensity of daily reproductive effort (*IDRE*), we used a modified version of the index developed by Burtschell et al. (2026) to accommodate for the absence of documented weaning mass in five species (Supporting Information 2). We used birth mass instead because it was highly correlated with weaning mass in Perissodactyla (Spearman’s rank correlation: *⍴* = 0.93, *p* = 0.01, *n* = 6 species tested):

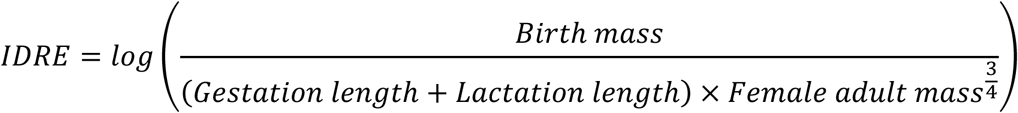

We retrieved the median social group size from Kamilar et al. (2010). Finally, the migratory behaviour was defined as a binary variable where 1 corresponds to migratory species, regardless of the distance and duration of the migration, whereas 0 corresponds to sedentary and nomadic species with consistent year-round home ranges (Webber and McGuire 2022).

### Predation exposure

As defining a predator as generalist or specialist towards a prey species is conceptually and methodologically challenging (e.g. it depends on the abundance of other prey species in a specific site, and may vary spatially and temporally), we designed a general predation exposure index based on two indices: the level of threat posed by a given predator species to a given prey species (*predation risk*) and the predator abundance in the study site of interest (*predator abundance*). Both indices ranged from 0 to 3 and corresponded to “absent”, “low”, “medium” and “high” (see details in Supporting Information 3). Although Perissodactyla are large prey relatively exempt from major predation risk except from large predators, juveniles of these species can be frequently preyed upon (Annear et al. 2023). In this analysis, we thus retained the main terrestrial predators present in the study sites: lion (*Panthera leo*), spotted hyena (*Crocuta crocuta*), leopard (*Panthera pardus*), wild dog (*Lycaon pictus*), cheetah (*Acinonyx jubatus*), tiger (*Panthera tigris*), snow leopard (*Panthera unica*), grey wolf (*Canis lupus*), and Nile crocodile (*Crocodylus niloticus*). We calculated the predation exposure index (*PEI*) as:

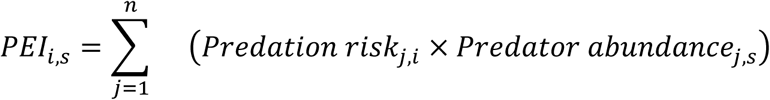

where *PEI_i,s_* is the predation exposure index of the prey species *i* in the study site *s*, *Predation risk_j,i_* is the index of selectivity of the predator species *j* toward the prey species *i*, and *Predator abundance_j,s_* is the index of abundance of the predator species *j* in the study site *s*. *PEI_i,s_* was an integer ranging between 0 and 27 in our dataset.

### Statistical analyses

We performed all analyses using the statistical software R (version 4.5.2; R Core Development Team, 2025) and stan (version 2.32.2; Stan Development Team, 2025). Predicted effects (*ꞵ*) are reported as the estimate ± standard error [95% credibility interval]). We also report 80% credibility intervals in the graphs.

We tested the effect of nine predictors (latitude, environmental seasonality, environmental productivity, environmental unpredictability, intensity of daily reproductive effort, migratory behaviour, pace of life, group size and predation exposure) on birth synchrony using Bayesian generalized linear mixed models with a Beta regression and a logit link function (package “brms”; Bürkner 2017). We standardized all continuous predictors by subtracting the mean and dividing by the standard deviation. Following Burtschell et al. (2026) recommendations, we first checked which of the dominant environmental predictors (productivity or seasonality) was most directly associated with the variation in reproductive seasonality using an additive bivariate model and found that environmental seasonality had the strongest explanatory power (see Results). Consequently, we used bivariate models including environmental seasonality as an additive control variable when testing the relationship between reproductive seasonality and predictors expected to show conditional effects with regards to environmental variability (environmental unpredictability, intensity of daily reproductive effort and migratory behaviour). We used univariate models for the other predictors. As there was no significant phylogenetic signal for reproductive seasonality (Blomberg’s *K* = 0.27, *p* = 0.11), we did not include phylogenetic relationship between species in the models. However, we included species identity as a random intercept to account for the presence of several populations of the same species. As sample sizes varied widely (*mean* ± *SD* = 232.8 ± 365.7; *range* = [12; 1555]) and can affect the robustness of birth synchrony estimates (Peláez et al. 2020), we used weighted regressions to account for the difference in sample sizes between populations, defined as: 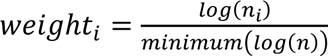, where *n_i_* is the sample size of the population *i* and *n* in the distribution of sample sizes of each population (Dezeure et al. 2024). The population with the smallest sample size thus represents one observation, while the other populations represent more observations depending on their sample size, following a logarithmic scale. For each model, we set an uninformative prior and used 3000 iterations, a burn-in of 1000 and three chains. We visually inspected for convergence and checked the absence of autocorrelations for the posterior distributions of fixed and random effects. The predictors were considered statistically significant when their 95% credibility intervals did not cross 0. We also investigated the correlation between the predictors using Spearman rank correlation tests (Supporting Information 4).

## Results

We retrieved birth data for 27 wild populations from 10 species (Table 1, Figure 1a), located between 33°31’S and 47°43’N (Figure 1b). *Equidae* accounted for 56% of the dataset and *Rhinocerotidae* accounted for 44%. We could not identify studies reporting the distribution of births for any *Tapiridae* species (*Tapirus terrestris*, *Tapirus bairdii*, *Tapirus indicus*, *Tapirus pinchaque*; but see Tortato et al. 2002 for a few observations in a semi-captive population), nor the African wild ass (but see Tesfai et al. 2021 for a general description) or the lesser one-horned rhinoceros (but see Setiawan and Yahya 2002 for a few observations). Birth synchrony varied between *r* = 0.06 for the greater one horned rhinoceros (*Rhinoceros unicornis*) in Chitwan National Park, Nepal, to 0.97 for the kiang in Chang Tang Reserve, China. Additionally, marked intra-specific variations were visible, particularly in the plains zebra (*range* = [0.24; 0.69], *n* = 7) and white rhinoceros (*Ceratotherium simum*, *range* = [0.07; 0.51 ], *n* = 3; Figure 1a, Fig. 2). Species identity was associated with significant random variance in all models.

**Figure 1:**
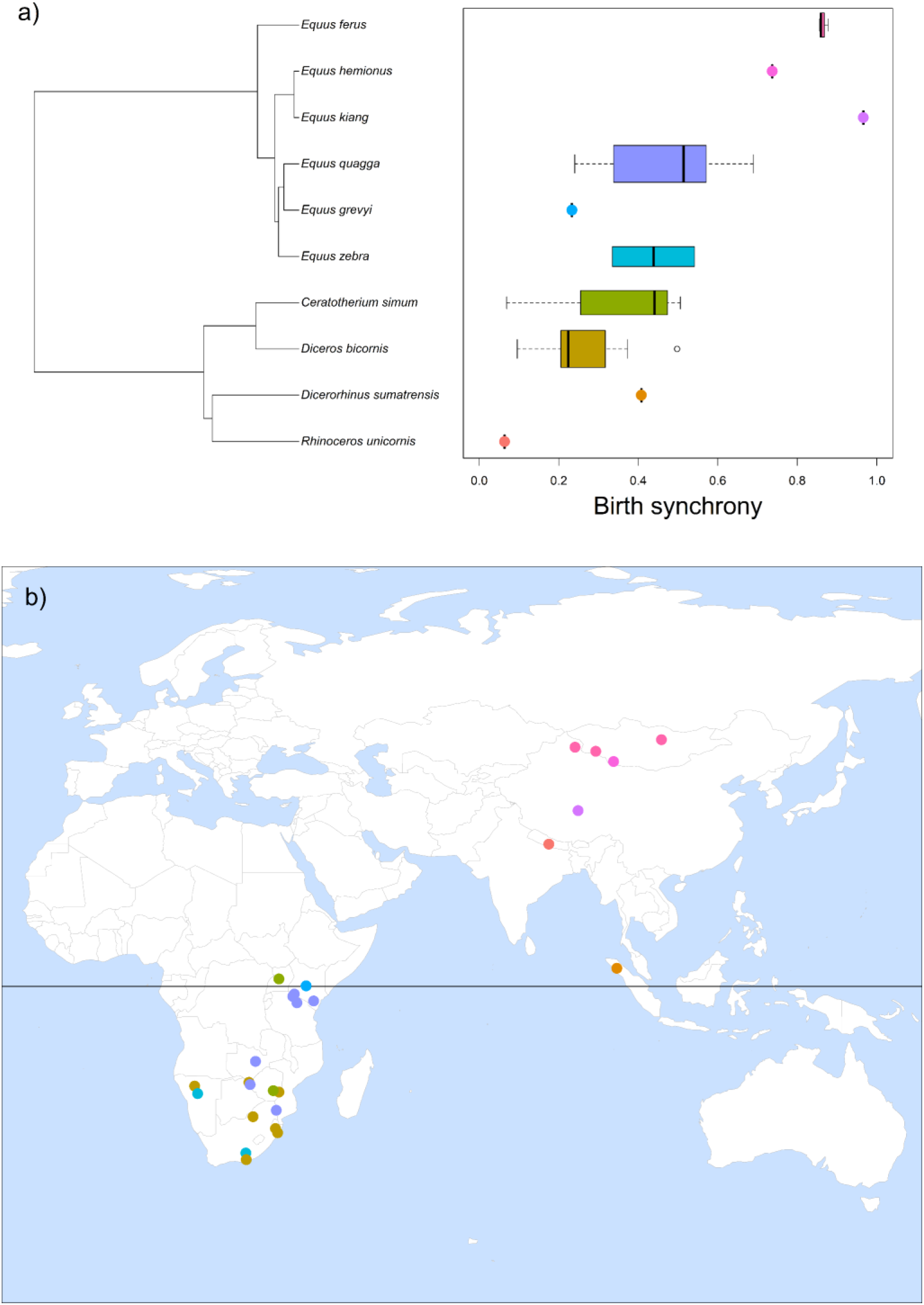
Populations of Perissodactyla (*n* = 27 populations, *n* = 10 species) retained in the analyses: a) population-specific distribution of birth synchrony (*r*) according to phylogenetic relatedness among species, b) location of the study sites (colours refer to species, as visible in panel a).

**Figure 2:**
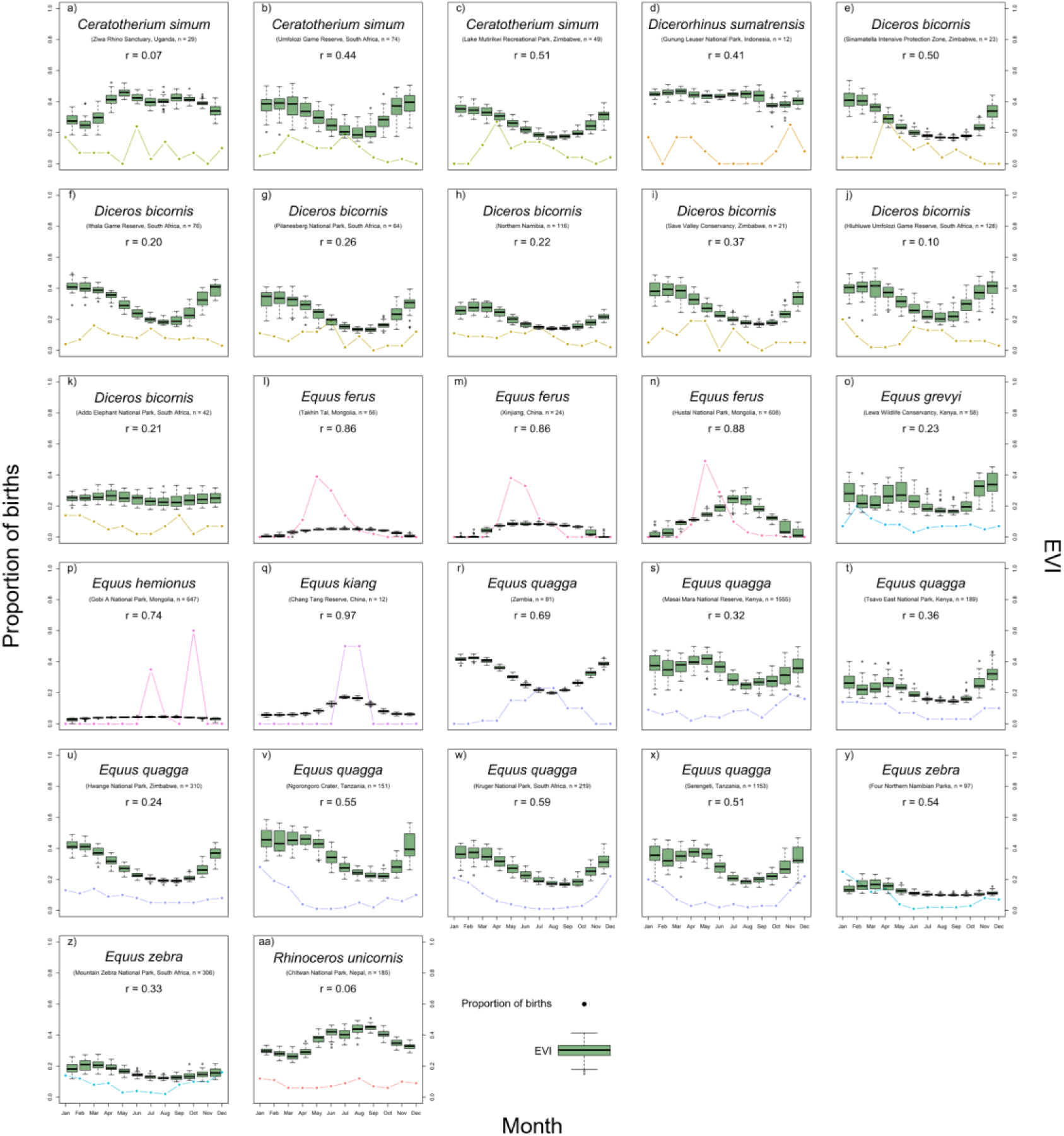
Monthly proportion of birth (solid dotted line) in relation to the monthly Enhanced Vegetation Index (EVI, boxplots) for each population (*n* = 27) of Perissodactyla species (*n* = 10). The EVI corresponds to the monthly averages between March 2000 and December 2024. Some study site names were shortened in the graph: “Four northern Namibian parks” stands for the combination of Etosha National Park, Daan Viljoen Game Reserve, Namib Desert Park and Naukluft Mountain Zebra Park, Namibia; “Save Valley Conservancy” stand for the combination of Save Valley Conservancy and Imire Game Ranch, Zimbabwe; “Hluhluwe Umfolozi Game Reserve” stands for Hluhluwe Corridor Umfolozi Game Reserve Complex, South Africa.

**Table 1:** Populations (*n* = 27) of Perissodactyla species (*n* = 10) included in the analyses. Study length is reported as the number of years during which birth dates were collected, “*r*” corresponds to the index of birth synchrony. Some study site names were shortened in the table: “Four northern Namibian parks” stands for the combination of Etosha National Park, Daan Viljoen Game Reserve, Namib Desert Park and Naukluft Mountain Zebra Park, Namibia; “Save Valley Conservancy” stand for the combination of Save Valley Conservancy and Imire Game Ranch, Zimbabwe; “Sinamatella IPZ” stands for Sinamatella Intensive Protection Zone, Zimbabwe; “Hluhluwe Umfolozi Game Reserve” stands for Hluhluwe Corridor Umfolozi Game Reserve Complex, South Africa.

| Species | Study site name | Study length | Sample size | $r$ |
| --- | --- | --- | --- | --- |
| *Ceratotherium simum* | Umfolozi Game Reserve, South Africa | 4 | 74 | 0.44 |
| *Ceratotherium simum* | Ziwa Rhino Sanctuary, Uganda | 11 | 29 | 0.07 |
| *Ceratotherium simum* | Lake Mutirikwi Recreational Park, Zimbabwe | 32 | 49 | 0.51 |
| *Dicerorhinus sumatrensis* | Gunung Leuser National Park, Indonesia | 6 | 12 | 0.41 |
| *Diceros bicornis* | Northern Namibia | 20 | 116 | 0.22 |
| *Diceros bicornis* | Addo Elephant National Park, South Africa | 26 | 42 | 0.21 |
| *Diceros bicornis* | Hluhluwe Umfolozi Game Reserve, South Africa | 13 | 128 | 0.10 |
| *Diceros bicornis* | Ithala Game Reserve, South Africa | 19 | 76 | 0.20 |
| *Diceros bicornis* | Pilanesberg National Park, South Africa | 17 | 64 | 0.26 |
| *Diceros bicornis* | Save Valley Conservancy, Zimbabwe | 6 | 21 | 0.37 |
| *Diceros bicornis* | Sinamatella IPZ, Zimbabwe | 5 | 23 | 0.50 |
| *Equus ferus* | Xinjiang, China | 5 | 24 | 0.86 |
| *Equus ferus* | Hustai National Park, Mongolia | 19 | 608 | 0.88 |
| *Equus ferus* | Takhin Tal, Mongolia | 8 | 56 | 0.86 |
| *Equus grevyi* | Lewa Wildlife Conservancy, Kenya | 5 | 58 | 0.23 |
| *Equus hemionus* | Gobi A National Park, Mongolia | 5 | 647 | 0.74 |
| *Equus kiang* | Chang Tang Reserve, China | 7 | 12 | 0.97 |
| *Equus quagga* | Masai Mara National Reserve, Kenya | 14.5 | 1555 | 0.32 |
| *Equus quagga* | Tsavo East National Park, Kenya | 2 | 189 | 0.36 |
| *Equus quagga* | Kruger National Park, South Africa | 4 | 219 | 0.59 |
| *Equus quagga* | Ngorongoro Crater, Tanzania | 3 | 151 | 0.55 |
| *Equus quagga* | Serengeti, Tanzania | 25 | 1153 | 0.51 |
| *Equus quagga* | Zambia | 2 | 81 | 0.69 |
| *Equus quagga* | Hwange National Park, Zimbabwe | 12 | 310 | 0.24 |
| *Equus zebra* | Four Northern Namibian Parks, Namibia | 3 | 97 | 0.54 |
| *Equus zebra* | Mountain Zebra National Park, South Africa | 30 | 306 | 0.33 |
| *Rhinoceros unicornis* | Chitwan National Park, Nepal | 15 | 185 | 0.06 |

Both the absolute latitude (*ꞵ_lati_* = 0.31 ± 0.17 [-0.03; 0.63], Figure 3a) and environmental seasonality (*ꞵ_seas_* = 0.76 ± 0.23 [0.31; 1.23], Figure 3b) were positively associated with birth synchrony in Perissodactyla. To the contrary, the environmental productivity was negatively associated with birth synchrony (*ꞵ_prod_* = -0.61 ± 0.20 [-1.00; - 0.19], Figure 3c). When both environmental seasonality and productivity were included in the same bivariate model, environmental seasonality (*ꞵ_seas_* = 0.58 ± 0.21 [0.18; 1.02]) had a stronger absolute coefficient relative to environmental productivity (*ꞵ_prod_* = -0.45 ± 0.20 [-0.83; -0.05]). There was no significant relationship between environmental unpredictability and birth synchrony, whether controlling for environmental seasonality (*ꞵ_unpr_* = -0.07 ± 0.12 [-0.30; 0.16], Figure 3d) or not (*ꞵ_unpr_* = -0.08 ± 0.13 [-0.33; 0.18]).

**Figure 3:**
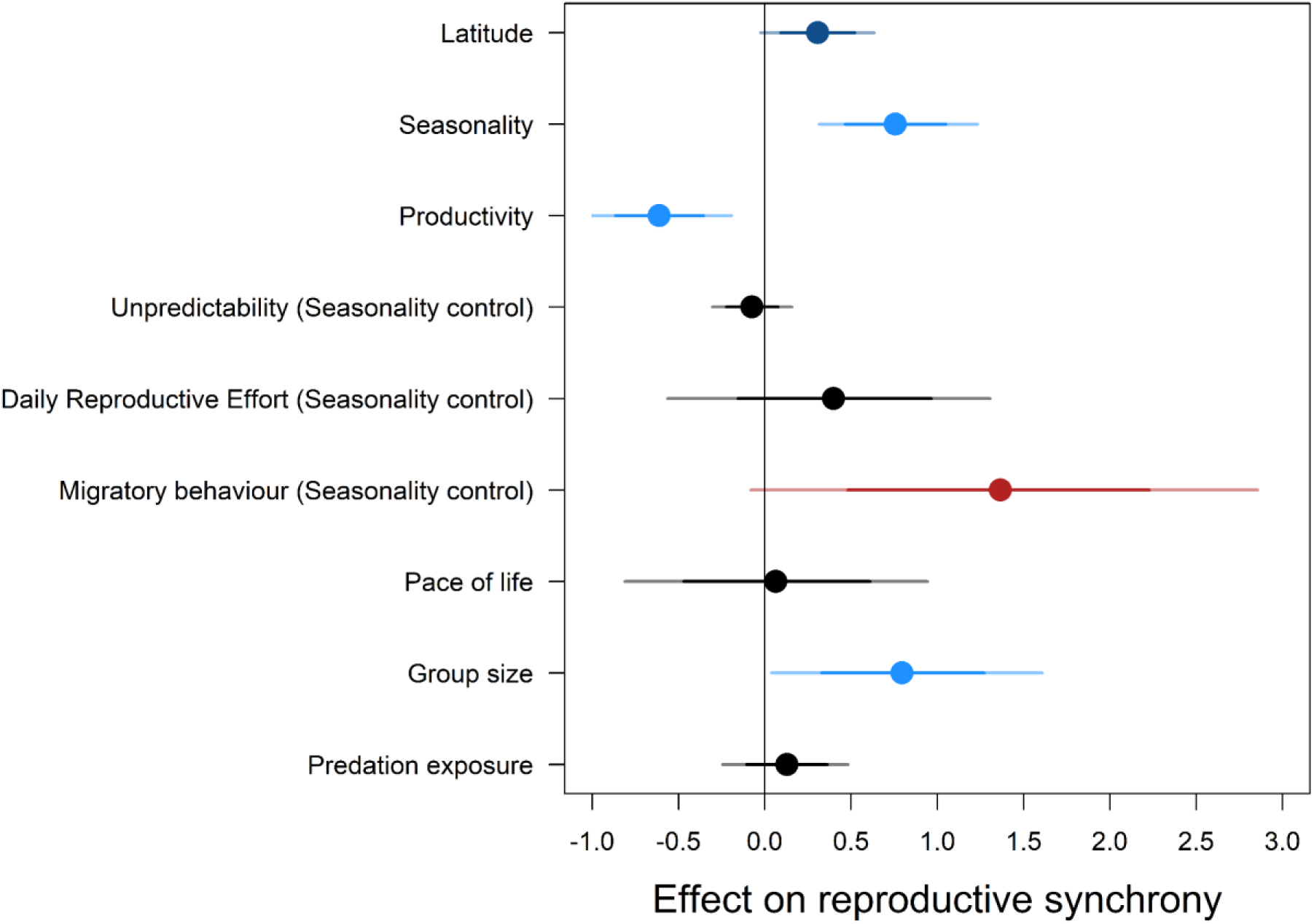
Effect of nine predictors on birth synchrony in 27 populations of Perissodactyla species (n = 10). Dots represent the mean effects estimated by the models (univariate models, unless specified otherwise: bivariate models with a control variable for environmental seasonality), light-coloured segments represent the 95% credibility interval of the mean effects, dark-coloured segments represent the 80% credibility interval of the mean effects. Shades of blue indicate a relationship in accordance with our predictions, shades of red indicate a relationship that is consistent with one of our alternative hypotheses, black indicates a non-significant relationship.

The intensity of daily reproductive effort had a non-significant positive effect on birth synchrony, whether controlling for environmental seasonality (*ꞵ_idre_* = 0.40 ± 0.46 [-0.56; 1.30], Figure 3e) or not (*ꞵ_idre_* = 0.78 ± 0.48 [-0.20; 1.74]). Although we found no significant effect of the migratory behaviour on birth synchrony, whether controlling for environmental seasonality (*ꞵ_migr_* = 1.37 ± 0.72 [-0.08; 2.86], Figure 3f) or not (*ꞵ_migr_* = 0.91 ± 1.09 [-1.25; 3.28]), migratory species marginally displayed a higher level of birth synchrony than non-migratory species. We found no significant effect of the pace of life on birth synchrony (*ꞵ_ppof_* = 0.06 ± 0.44 [-0.81; 0.94], Figure 3g).

Finally, group size had a significant positive effect on birth synchrony (*ꞵ_gsiz_* = 0.80 ± 0.39 [0.04; 1.61], Figure 3h), but predation exposure did not significantly affect birth synchrony (*ꞵ_pred_* = 0.13 ± 0.18 [-0.24; 0.48], Figure 3i).

## Discussion

Using a comparative analysis, we investigated the main determinants of synchrony of birth in 10 Perissodactyla species. As expected, we found a strong effect of environmental predictors with an increase in birth synchrony with environmental seasonality (and to a lower extent latitude), and a decrease in birth synchrony with environmental productivity, but we found no effect of environmental unpredictability. Life history traits did not have a strong effect on birth synchrony among Perissodactyla: the intensity of daily reproductive effort and the pace of life did not affect birth synchrony significantly, but migratory behaviour marginally increased birth synchrony, as it allowed access to a broader peak of seasonal resources, rather than buffering resource variation. In line with our predictions, group size increased birth synchrony, but the effect of predation exposure was not significant.

We found a strong positive correlation between the level of birth synchrony and environmental seasonality, and a strong negative correlation with environmental productivity, both predictors being negatively correlated in our dataset (Supporting information 4). The marginally positive effect of absolute latitude also confirms an already extended literature among large herbivores (e.g. Rutberg 1987, English et al. 2012, Zerbe et al. 2012), but also numerous other mammal groups (e.g. in carnivores, Heldstab et al. 2018; in lagomorphs, Heldstab 2021a; in primates, Heldstab et al. 2021; in rodents, Heldstab 2021b). However, we found no strong support for a correlation between environmental unpredictability and birth asynchrony, contrary to similar comparative analyses (in large herbivores, English et al. 2012; in primates, Dezeure et al. 2024, Burtschell et al. 2026). Our absence of significant results might come from the small variability in terms of environmental unpredictability among the study sites inhabited by the species included in this study, most of them being characterised by fairly predictable environments (Figure 4d).

**Figure 4:**
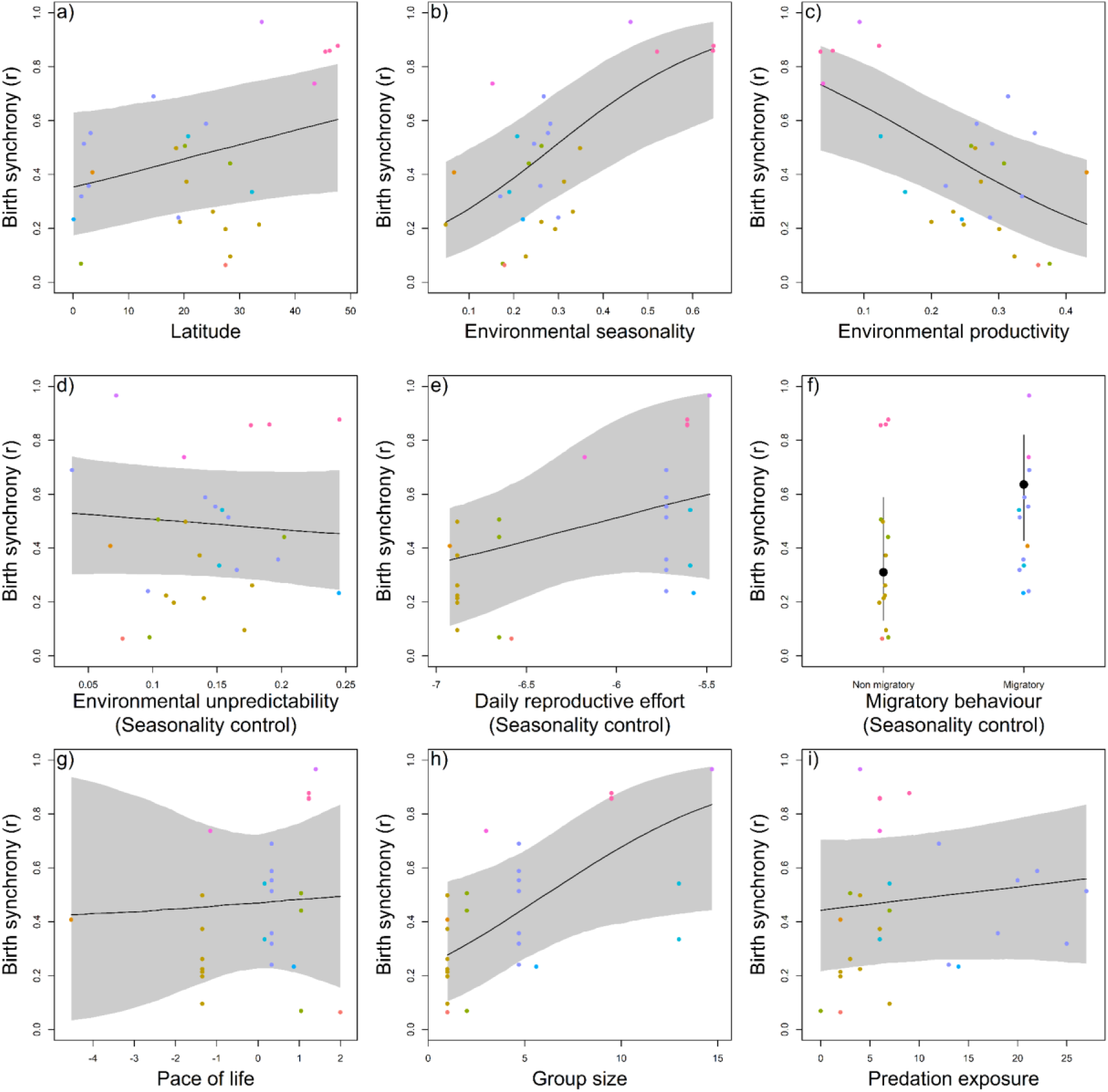
Predicted effect (for an averaged environmental seasonality as a control variable when appropriate, see text for detail) of nine predictors on birth synchrony in 27 populations of Perissodactyla species (*n* = 10): a) latitude, b) environmental seasonality, c) environmental productivity, d) environmental unpredictability, e) intensity of daily reproductive effort, f) migratory behaviour, g) pace of life, h) group size, i) predation exposure. Dots represent observed values for each population (colours indicating the corresponding species), the solid line represents the predictions of the models (for an averaged environmental seasonality as a control variable, when appropriate, see text for detail), the shaded area (black segments, when appropriate) represents the 95% credibility interval.

Although the species played a significant role in shaping the relationships between the nine predictors and birth synchrony in our populations, we found no effect of phylogenetic relatedness on birth synchrony, and a strong inter-population variability was visible (Figure 1a, Figure 2). Similar population effects have been described in Artiodactyla species within their natural range, such as the impala (*Aepyceros melampus*, Moe et al. 2007) or the roe deer (*Capreolus capreolus*, Peláez et al. 2020). Ten of the 11 studied species exhibit lower synchrony in captivity, where food resources are accessible *ad libitum* and predation absent (Zerbe et al. 2012). Numerous species are adjusting their reproductive timing in response to climate change (e.g. Renaud et al. 2019, Bonnet et al. 2019). Together, these results suggest that, additionally to ancestral evolutionary drivers, these two ecologically similar groups (Artiodactyla and Perissodactyla) can display a high flexibility and adjust their birth synchrony to pressures imposed by the current conditions experienced by the individuals. Although these patterns are often suggested to be primarily driven by environmental effects, the role of other factors such as predation cannot be excluded (Moe et al. 2007, Peláez et al. 2020).

Nevertheless, we found limited evidence for a positive relationship between birth synchrony and our predation exposure index, the effect being non-significant. This lack of significance may reflect the fact that predation acts as a second-order pressure on phenology patterns, after environmental factors (Zerbe et al. 2012). Additionally, our predation exposure index does not account for site-specific selectivity of a given predator species for a given prey species, which also depends on the other prey species available to the predator as well as the other predator species competing over prey access in that specific site (Ferretti et al. 2020). Consequently, our generalist approach might have diluted the strength of the relationship between predation exposure and birth synchrony.

More generally, studying the effect of predation on birth synchrony remains a challenging task, both methodologically (difficulty to collect the adequate data for sound empirical testing) and conceptually: several factors can interact such as the anti-predator strategy of the juvenile and group size, both generally correlated and influencing birth synchrony (this study, Sinclair et al. 2000). Both *Equidae* (Wilson and Mittermeier 2011) and *Rhinocerotidae* (e.g. Medhi and Sakia 2019) are predominantly classified as follower species (i.e. the juvenile follows the mother soon after birth, Lent 1974), and our results conform with the theoretical expectation that among followers, the more they are exposed to specialist predators, the more synchronized their births should be (Rutberg 1987). However, other empirical examples among hider species (which hide their juveniles for a few days to a few weeks after birth) suggest that contrary to theoretical expectations, the best anti-predator strategy against generalist predators might also be to synchronise births (Michel et al. 2020). Including hider species such as the *Tapiridae* (Wilson and Mittermeier 2011) and their guild of predators might provide more robust conclusions.

Migratory species tended to have more synchronous births than non-migratory species. Migration could act as a temporal constraint on reproduction, individuals having to trade off the timing of birth with the timing of migration depending on their migratory strategy (Bischof et al. 2012, Aikens et al. 2021). However, Perissodactyla are mostly classified as capital breeders (Thel et al. 2025) and should rely more on long-term storage than readily available food resources for their reproduction, thus diluting the direct link between the timing of food resource availability and reproductive cycle, to the point that migration and parturition can be completely decorrelated (e.g. Laforge et al. 2023). Additionally, most of the migratory species included in our analyses being found in productive subtropical environments, migration in these ecosystems might be a strategy to capitalise on spatialised productivity peaks rather than to escape starvation. Similarly to behavioural buffers in primates (Burtschell et al. 2026), migration could consequently not act as a driver of reproductive phenology, but rather as a covariate in these ecosystems, both migration and reproduction temporally and spatially synchronising with the most favourable conditions. Migratory behaviour was also correlated with other non-environmental factors such as the intensity of daily reproductive effort and predation exposure in our dataset (Supporting information 4). In the highly migratory blue wildebeest, the high level of birth synchrony has been attributed to coinciding anti-predator strategies more than the exposure to environmental constraints (Estes 1976).

Ultimately, classifying migratory behaviours can be challenging as some species have migratory and non-migratory populations, this behaviour even varying in a given individual during its lifetime (Estes 1976, Thel et al. 2025). With our general approach, we also could not quantify how migration modifies the environment experienced by a population and if it actually reduces the seasonality they are exposed to in terms of environment and food resources. Overall, the link between migratory behaviour and birth phenology is not direct and requires further theoretical development and empirical exploration to be conclusively investigated.

We found no significant relationship between the pace of life and birth synchrony. Reproductive traits tend to be highly correlated with body mass (e.g. gestation length in large herbivores, Owen-Smith and Ogutu 2013), but covariation between life history traits can persist even after controlling for body mass, encapsulated in the concept of pace of life (Bielby et al. 2007, Healy et al. 2019). However, we found that Perissodactyla display fairly homogeneous paces of life, the only species having an unexpectedly slow pace of life relative to its body mass being the Sumatran rhinoceros (Supporting information 2). Such homogeneity among this order limited our ability to test for an effect of pace of life and suggests that pace of life might not be an influential factor in the variability of birth synchrony among Perissodactyla. In general, as expected for a lineage with generally slow pace of life, seasonality among most populations in our sample was low (average *r* below 0.50). However, our results suggest a positive trend between the intensity of the daily reproductive effort and birth synchrony, despite being non-significant. The less concentrated in time the energetic investment is, the less constrained the birth timing needs to be. Similar results were found in another group of slow living species, namely primates, which confirms the primary importance of life history, and specifically the intensity of daily reproductive effort, in shaping reproductive phenology (Burtschell et al. 2023, 2026).

Finally, we found no study on the distribution of births in *Tapiridae*, nor in the lesser one-horned rhinoceros (Wilson and Mittermeier 2011). These solitary, elusive and mostly nocturnal species (e.g. Hariyadi et al. 2010, Cruz et al. 2014) inhabit difficult to access locations, where population monitoring is challenging, probably explaining why their reproduction in the wild has been understudied so far (Thel et al. 2026). We made a similar observation for the African wild ass, a species with a very limited geographical range. Most of these species being currently endangered to critically endangered, we encourage scientists to use and share by-catch data (e.g. from large-scale camera trap monitoring) and develop non-invasive population monitoring programs to expand the body of knowledge on the reproduction of these species.

## Conclusion

Our study confirmed the dominant role of environmental variables (principally seasonality, productivity and latitude) in shaping the distribution of births in Perissodactyla, a currently under-studied order among large herbivores, at the global scale. It also suggests more robust empirical testing is required to confirm the positive effect of predation and migration on birth synchrony, as the quantitative assessment of such factors and their correlated nature (with each other and with other environmental and life history traits) makes their effects difficult to disentangle. While the exact factors shaping reproductive timing require additional study, our findings do indicate that Perissodactyla shift their breeding in response to local conditions, leading to local adaptation even among populations of the same species, highlighting the potential risk of mismatches if individuals are moved to new sites.

## Data availability statement

Data available from the Zenodo Repository: https://doi.org/10.5281/zenodo.20586291.

## Authors’ contributions

LT: Conceptualization - Methodology - Validation - Formal analysis - Investigation - Data Curation - Writing: Original Draft - Writing: Review & Editing - Visualisation - Funding acquisition

EH: Conceptualization - Methodology - Writing: Review & Editing

DL: Formal analysis - Methodology - Writing: Review & Editing

BG: Conceptualization - Writing: Review & Editing

JD: Methodology - Writing: Review & Editing

JAV: Conceptualization - Methodology - Writing: Review & Editing - Funding acquisition

## Acknowledgments

We thank Lugdiwine Burtschell for her suggestions during the early-stage conceptualization of the project. This research was funded by a Postgraduate Research Scholarship from Nelson Mandela University attributed to LT.

