## supporting information for "Effect of the environment, predation and life history on reproductive synchrony in Perissodactyla"

Supporting information 1: General information on the populations of Perissodactyla retained in the study.

### *Data selection*

Several populations of zebra and African rhinoceros living in fully fenced reserves (*n* = 9) or in semi-captive condition (*n* = 1) were kept in the analyses as most of the wildlife is kept in fenced reserves in Eastern and Southern Africa (Packer et al. 2013, Pekor et al. 2019) and most rhinoceros populations nowadays are closely managed due to heavy poaching threats (Clements et al. 2020). Excluding these studies would have severely reduced the sample size and although these populations currently occur in modified ecosystems (e.g. modified predator densities), we assumed here that fences were erected too recently to interact with birth phenology at the evolutionary scale in long-lived species such as Perissodactyla (Hayward et al. 2009). Additionally, we used predictors describing the current state of the environments experienced by the populations of interest, thus accounting for such deviation from fully natural systems (see Methods section in the main text). We did not include populations outside of their historical range (e.g. white rhinoceros in the Eastern Cape, South Africa, Truter 2021).

No quantitative description of the phenology of births was available for the kiang, but as this species was recurrently described as a highly seasonal breeder foaling in July and August (Schaller 1998, Sharma et al. 2004, Paklina and van Orden 2007, St-Louis and Côté 2009), we generated an artificial distribution of births to include this species in our analyses. We attributed the smallest number of births for a study to be included in the analyses (*n* = 12 births) and spread them evenly between July and August for the study site where the monitoring was >1 year (Chang Tang Reserve, China, Schaller 1998).

***Data preparation***

When studies only reported bi-monthly counts, we divided the number of births equally between the two months ($n$ = 3 studies). If several studies described birth phenology for the same population (i.e. same species and same study site) but the period of data collection differed and did not overlap, we added both distributions to increase sample size (n = 5 populations).

Table S1.1: Details about the population, study site, data collection and data processing (both from the original study and from our data processing for inclusion in our analyses) for the populations of Perissodactyla included in the analyses ($n$ = 27).

| **Population reference code** | **Species scientific name (reference)** | **Species scientific name (in the study, if different)** | **Study site name (reference)** | **Study site name (in the study, if different)** | **Study site status** | **Study site coordinates note** | **Study length** | **Sample size** | **Sample size notes** | **Birth phenology notes** | **Birth phenology source (in the original study)** | **Combined multiple studies** | **Reference** | **Type of reference** | **General remarks** |
|---|---|---|---|---|---|---|---|---|---|---|---|---|---|---|---|
| DbP | *Diceros bicornis* | | Pilanesberg National Park, South Africa | | fully fenced | from latitude.to (error in the study) | 17 | 64 | sample size different between the graph and the text, sample size from the graph retained | conceptions dates reported, but estimated from the dates of birth, used to back calculate dates of birth | Fig.3a | no | Hrabar and Du Toit 2005 | peer reviewed article | none |
| EqT | *Equus quagga* | *Equus burchelli bohmi* | Tsavo East National Park, Kenya | | open | from latitude.to (coordinates not provided) | 2 | 189 | sample size calculated: use percentage of young foals relative to the number of adults, which ranges between 167 and 729 between surveys, averaged to 448 | bi-monthly distribution spread equally between both months | Fig. 2 | no | Leuthold and Leuthold 1975 | peer reviewed article | none |
| EqZ | *Equus quagga* | *Equus burchelli* | Zambia | Northern Rhodesia | open | from latitude.to (barycentre of the country) | 2 | 81 | none | 1) bi-monthly distribution spread equally between both months. 2) no scale on the figure: proportions estimated based on the total sample size | Fig. 1 | no | Ansell 1960 | peer reviewed article | none |
| CsL | *Ceratotherium simum* | *Ceratotherium simum simum* | Lake Mutirikwi Recreational Park, Zimbabwe | Kyle National Park | fully fenced | from the study | 10 | 15 | none | none | Fig. 5 | yes (Condy 1973: from 1962 to 1972, Monks 1995: from 1973 to 1995) | Condy 1973 | master's thesis | none |
| CsL | *Ceratotherium simum* | *Ceratotherium simum simum* | Lake Mutirikwi Recreational Park, Zimbabwe | Kyle Recreational Park | fully fenced | from the study | 22 | 34 | sample size different between the graph and the text, sample size from the graph retained | conception and birth on the same graph with same legend: we consider conception on the left and birth on the right according to description of mating and calving periods in the text | Fig. 4 | yes (Condy 1973: from 1962 to 1972, Monks 1995: from 1973 to 1995) | Monks 1995 | master's thesis | none |
| RuC | *Rhinoceros unicornis* | | Chitwan National Park, Nepal | Royal Chitawan National Park | open | from Dinerstein and Price 1991 | 3 | 60 | none | 1) bi-monthly distribution spread equally between both months. 2) counts from two study sites added: Sauraha and Tiger Tops approx. 25 km apart from each other | Tab. 3.14 | yes (Laurie 1978: from 1972 to 1975, Dinerstein and Price 1991: from 1984 to 1988, Subedi et al. 2017: from 2009 to 2015) | Laurie 1978 | PhD thesis | none |
| RuC | *Rhinoceros unicornis* | | Chitwan National Park, Nepal | Royal Chitwan National Park | open | from the study | 5 | 59 | sample size different between the graph and the text, sample size | none | Fig. 2 | yes (Laurie 1978: from 1972 to 1975, Dinerstein and Price 1991: from 1984 to | Dinerstein and Price 1991 | peer reviewed article | none |

| Population reference code | Species scientific name (reference) | Species scientific name (in the study, if different) | Study site name (reference) | Study site name (in the study, if different) | Study site status | Study site coordinates note | Study length | Sample size | Sample size notes | Birth phenology notes | Birth phenology source (in the original study) | Combined multiple studies | Reference | Type of reference | General remarks |
|---|---|---|---|---|---|---|---|---|---|---|---|---|---|---|---|
| | | | | | | | | | from the graph retained | | | 1988, Subedi et al. 2017: from 2009 to 2015) | | | |
| RuC | *Rhinoceros unicornis* | | Chitwan National Park, Nepal | | open | from Dinerstein and Price 1991 | 7 | 66 | none | number of calves not integers | Fig. 1 | yes (Laurie 1978: from 1972 to 1975, Dinerstein and Price 1991: from 1984 to 1988, Subedi et al. 2017: from 2009 to 2015) | Subedi et al. 2017 | peer reviewed article | none |
| EqK | *Equus quagga* | *Equus (Hippotigris) burchelli antiquorum* | Kruger National Park, South Africa | | open | from the study | NA | 39 | none | none | Fig. 19 | yes (Fairall 1968: up to 1968, Smuts 1976: from 1969 to 1973) | Fairall 1968 | peer reviewed article | none |
| EqK | *Equus quagga* | *Equus burchelli antiquorum* | Kruger National Park, South Africa | | open | from Fairall 1968 | 4 | 180 | no sample size of foals provided, sample size of embryos taken instead | none | Tab. 6 | yes (Fairall 1968: up to 1968, Smuts 1976: from 1969 to 1973) | Smuts 1976 | peer reviewed article | none |
| DbA | *Diceros bicornis* | *Diceros bicornis michaeli* | Addo Elephant National Park, South Africa | | fully fenced | from Freeman et al. 2014 | 15 | 12 | none | none | Fig. 1 | yes (Hall-Martin and Penzhorn 1977: from 1961 and 1975, Freeman et al. 2014: from 2001 to 2012) | Hall-Martin and Penzhorn 1977 | peer reviewed article | authors note: individuals translocated from Kiboko, Kenya and subject to social conditions which they would not normally have encountered in the wild |
| DbA | *Diceros bicornis* | *Diceros bicornis bicornis* | Addo Elephant National Park, South Africa | | fully fenced | from the study | 11 | 30 | total of the percentages is < 100 | counts from two study sites added: Addo and Nyathi, next to each other | Fig. 2 | yes (Hall-Martin and Penzhorn 1977: from 1961 and 1975, Freeman et al. 2014: from 2001 to 2012) | Freeman et al. 2014 | peer reviewed article | none |
| EqS | *Equus quagga* | | Serengeti, Tanzania | | open | from latitude.to (coordinates not provided) | 4 | 705 | none | none | Fig. 6b | yes (Klingel 1969: from 1962 to 1965, Sinclair et al. 2000: from 1977 to 1997) | Klingel 1969 | peer reviewed article | same population and data collection period as Klingel 1965 |
| EqS | *Equus quagga* | *Equus burchelli* | Serengeti, Tanzania | | open | from latitude.to (coordinates not provided) | 21 | 448 | none | use counts but percentages also provided in the study | Tab. 2 | yes (Klingel 1969: from 1962 to 1965, Sinclair et al. 2000: from 1977 to 1997) | Sinclair et al. 2000 | peer reviewed article | none |
| EkC | *Equus kiang* | | Chang Tang Reserve, China | | open | from the study | 7 | 12 | sample size generated artificially | birth distribution generated from qualitative description | generated artificially | no | Schaller 1998 | book | none |
| DbS | *Diceros bicornis* | | Sinamatella Intensive Protection Zone, Zimbabwe | | open | from latitude.to (coordinates not provided) | 5 | 23 | none | none | Tab. 3 | no | Alibhai et al. 2001 | peer reviewed article | 1) Sinamatella Intensive Protection Zone located in Hwange National Park. 2) some |

| Population reference code | Species scientific name (reference) | Species scientific name (in the study, if different) | Study site name (reference) | Study site name (in the study, if different) | Study site status | Study site coordinates note | Study length | Sample size | Sample size notes | Birth phenology notes | Birth phenology source (in the original study) | Combined multiple studies | Reference | Type of reference | General remarks |
|---|---|---|---|---|---|---|---|---|---|---|---|---|---|---|---|
| | | | | | | | | | | | | | | | females immobilised |
| EfH | *Equus ferus* | *Equus przewalskii* | Hustai National Park, Mongolia | | open | from the study | 19 | 608 | none | none | Fig. 2 | no | Dorj and Namkhai 2013 | peer reviewed article | none |
| EhG | *Equus hemionus* | | Gobi A National Park, Mongolia | Gobi B National Park, but does not correspond to the current maps, corresponds to Gobi A instead | open | from latitude.to (coordinates not provided) | 5 | 647 | none | none | Tab. 6 | no | Feh et al. 2001 | peer reviewed article | 1992 excluded because data collected mostly before the birth peak |
| DbV | *Diceros bicornis* | *Diceros bicornis minor* | Save Valley Conservancy and Imire Game Ranch, Zimbabwe | | some study sites fully fenced (Imre Game Ranch) | Save Valley Conservancy, from latitude.to (error in the coordinates) | 6 | 21 | none | none | Fig. 5 | no | Garnier et al. 2002 | peer reviewed article | same population and data collection period as Garnier 2001 |
| DbI | *Diceros bicornis* | | Ithala Game Reserve, South Africa | | fully fenced | from the study | 19 | 76 | none | none | Fig. 3 | no | Greaver et al. 2014 | peer reviewed article | none |
| EqE | *Equus zebra* | *Equus zebra hartmannae* | Etosha National Park and Daan Viljoen Game Reserve and Namib Desert Park and Naukluft Mountain Zebra Park, Namibia | South West Africa | some study sites fully fenced (Etosha National Park and Daan Viljoen Game Reserve) | averaged between the four study sites, from latitude.to | 3 | 97 | none | none | Fig. 3 | no | Joubert 1974 | peer reviewed article | study sites and study period extracted from Joubert 1972 |
| EqN | *Equus quagga* | *Equus quagga boehmi* | Ngorongoro Crater, Tanzania | | open | from latitude.to (coordinates not provided) | 3 | 151 | none | none | Fig. 1 | no | Klingel 1965 | peer reviewed article | same population and data collection period as Klingel 1969 |
| DbN | *Diceros bicornis* | | North, Namibia | | open | Ongava farm (provided the permits), from latitude.to (coordinates not provided) | 20 | 116 | none | none | Fig. 3a | no | Muntifering et al. 2023 | peer reviewed article | none |
| EqM | *Equus quagga* | *Equus burchelli* | Masai Mara National Reserve, Kenya | | open | from the study | 14.5 | 1555 | none | none | Tab. 2 | no | Ogutu et al. 2013 | peer reviewed article | same population and data collection period as Ogutu et al. 2011 and Ogutu et al. 2014b |
| CsU | *Ceratotherium simum* | | Umfolozi Game Reserve, South Africa | | fully fenced | from latitude.to (coordinates not provided) | 4 | 74 | none | none | Fig. 7.7 | no | Owen-Smith 1988 | book | none |
| CsZ | *Ceratotherium simum* | | Ziwa Rhino Sanctuary, Uganda | | fully fenced | from latitude.to (coordinates not provided) | 11 | 29 | none | none | Tab. C.2.5.1 | no | Patton and Genade 2021 | technical report | 1) same population and data collection period as Patton et al. 2022. 2) highly modified ecosystem |

| Population reference code | Species scientific name (reference) | Species scientific name (in the study, if different) | Study site name (reference) | Study site name (in the study, if different) | Study site status | Study site coordinates note | Study length | Sample size | Sample size notes | Birth phenology notes | Birth phenology source (in the original study) | Combined multiple studies | Reference | Type of reference | General remarks |
|---|---|---|---|---|---|---|---|---|---|---|---|---|---|---|---|
| EzM | *Equus zebra* | *Equus zebra zebra* | Mountain Zebra National Park, South Africa | | fully fenced | from the study | 30 | 306 | none | none | Fig. 3 | no | Penzhorn 1985 | peer reviewed article | none |
| EqH | *Equus quagga* | | Hwange National Park, Zimbabwe | | open | from the study | 12 | 310 | none | none | Fig. 1 | no | Thel et al. 2025 | peer reviewed article | none |
| EfX | *Equus ferus* | *Equus przewalskii* | Xinjiang, China | | open | from the study | 5 | 24 | none | Fig. 3 provides slightly different counts | Fig. 2 | no | Chen et al. 2008 | peer reviewed article | none |
| DsG | *Dicerorhinus sumatrensis* | | Gunung Leuser National Park, Indonesia | | open | from the study | 6 | 12 | none | date of birth estimated based on the size of footmarks: up to 7-month uncertainty | Fig. 6.1 | no | Van Strien 1985 | PhD thesis | none |
| EgL | *Equus grevyi* | | Lewa Wildlife Conservancy, Kenya | | fully fenced | from the study | 5 | 58 | sample size rounded to the nearest integer | none | Fig. 1.2 | no | Tupper 2011 | master's thesis | none |
| DbH | *Diceros bicornis* | *Diceros bicornis minor* | Hluhluwe Corridor Umfolozi Game Reserve Complex, South Africa | | fully fenced | from latitude.to (coordinates not provided) | 13 | 128 | none | none | Fig. 1 | no | Hitchins and Anderson 1983 | peer reviewed article | none |
| EfT | *Equus ferus* | *Equus ferus przewalskii* | Takhin Tal, Mongolia | | open | from latitude.to (coordinates not provided) | 8 | 56 | none | only the birth after 1999 were retained to exclude captive-born individuals | Tab. 1 | no | Hoesli et al. 2009 | non-peer reviewed article | none |

Study length is reported as the number of years during which birth dates were collected.

Supporting information 2: Species-specific life history traits.

Social group size was determined as the median in the reference study (Kamilar et al. 2010). For species not reported by Kamilar et al. (2010), we used other references and calculated the median group size when multiple values were reported to match the reference study. Female body mass was extracted from Myhrvold et al. (2015). For species not reported by Myhrvold et al. (2015), we used other references and calculated the mean female body mass when multiple values were reported. We could not identify a reference providing female body mass for the onager, so we used adult body mass instead (Myhrvold et al. 2015). Migratory behaviour was extracted from Webber and McGuire (2022), and defined as a binary variable where 1 corresponds to migratory species, regardless of the distance and duration of the migration, whereas 0 corresponds to sedentary and nomadic species with consistent year-round home ranges (Webber and McGuire 2022). For species not reported by Webber and McGuire (2022), we used other references and followed the same definition as the reference study.

Table S2.1: Species-specific life history traits collected for the species of Perissodactyla included in this study (*n* = 10).

| **Species scientific name (reference)** | **Social group size** | **Social group size (reference)** | **Migratory behaviour** | **Migratory behaviour (reference)** | **Birth mass** | **Birth mass (reference)** | **Weaning mass** | **Weaning mass (reference)** | **Female mass** | **Female mass (reference)** | **Gestation length** | **Gestation length (reference)** | **Lactation length** | **Lactation length (reference)** | **Interbirth interval** | **Interbirth interval (reference)** | **Female age at maturity** | **Female age at maturity (reference)** |
|---|---|---|---|---|---|---|---|---|---|---|---|---|---|---|---|---|---|---|
| *Ceratotherium simum* | 2 | Owen-Smith 1988 | 0 | Webber and McGuire 2022 | 51167 | Myhrvold et al. 2015 | 274482 | Myhrvold et al. 2015 | 1600000 | Owen-Smith 1988 | 515 | Myhrvold et al. 2015 | 365 | Myhrvold et al. 2015 | 987 | Myhrvold et al. 2015 | 2026 | Myhrvold et al. 2015 |
| *Dicerorhinus sumatrensis* | 1 | Hutchins and Kreger 2006 | 1 | Webber and McGuire 2022 | 23000 | Myhrvold et al. 2015 | NA | / | 690000 | Roth et al. 2013 (from captive individuals) | 475 | Roth et al. 2004 (from captive individuals) | 502 | Myhrvold et al. 2015 | 1277 | Myhrvold et al. 2015 | 2739 | Myhrvold et al. 2015 |
| *Diceros bicornis* | 1 | Kamilar et al. 2010 | 0 | Webber and McGuire 2022 | 35000 | Myhrvold et al. 2015 | 426865 | Myhrvold et al. 2015 | 1000000 | Owen-Smith 1988 | 474 | Myhrvold et al. 2015 | 606 | Myhrvold et al. 2015 | 1053 | Myhrvold et al. 2015 | 1826 | Myhrvold et al. 2015 |
| *Equus ferus* | 9.5 | Hoesli et al. 2009 | 0 | King 2012 | 30000 | https://ielc.libguides.com/sdzg/factsheets/przewalskishorse/summary | NA | / | 280000 | Kuntz et al. 2006 (from semi-captive individuals) | 336 | Boyd and Houpt 1994 | 335 | Boyd 1991 (from captive individuals) | 365 | https://ielc.libguides.com/sdzg/factsheets/przewalskishorse/reproduction | 1095 | Rödel et al. 2023 (from semi-captive individuals) |
| *Equus grevyi* | 5.6 | Williams 1998, Sundaresan et al. 2007 | 1 | Webber and McGuire 2022 | 40000 | Myhrvold et al. 2015 | NA | / | 385000 | Tupper 2011 | 406 | Myhrvold et al. 2015 | 275 | Myhrvold et al. 2015 | 575 | Myhrvold et al. 2015 | 1342 | Myhrvold et al. 2015 |
| *Equus hemionus* | 3 | Kamilar et al. 2010, Sundaresan et al. 2007, Nowzari et al. 2013 | 1 | Lushchekina et al. 2022 | 17500 | Hering-Hagenbeck and Prahl (from captive individuals) | NA | / | 230000 | Myhrvold et al. 2015 (female weight unavailable, adult body mass retained instead) | 346 | Myhrvold et al. 2015 | 456 | Myhrvold et al. 2015 | 502 | Myhrvold et al. 2015 | 1157 | Myhrvold et al. 2015 |
| *Equus kiang* | 14.7 | Schaller 1998, St-Louis and Côté 2009 | 1 | St-Louis and Côté 2009 | 36000 | Myhrvold et al. 2015 | NA | / | 275000 | Myhrvold et al. 2015 | 304 | Myhrvold et al. 2015 | 418 | Myhrvold et al. 2015 | 547 | Myhrvold et al. 2015 | 1001 | Myhrvold et al. 2015 |
| *Equus quagga* | 4.7 | Klingel 1969 | 1 | NA | 32000 | Myhrvold et al. 2015 | 205000 | Myhrvold et al. 2015 | 302000 | Myhrvold et al. 2015 | 365 | Myhrvold et al. 2015 | 395 | Myhrvold et al. 2015 | 643 | Myhrvold et al. 2015 | 907 | Myhrvold et al. 2015 |
| *Equus zebra* | 13 | Kamilar et al. 2010 | 1 | Webber and McGuire 2022 | 30000 | Myhrvold et al. 2015 | 117098 | Myhrvold et al. 2015 | 276300 | Joubert 1974 | 363 | Myhrvold et al. 2015 | 304 | Myhrvold et al. 2015 | 662 | Myhrvold et al. 2015 | 1009 | Myhrvold et al. 2015 |
| *Rhinoceros unicornis* | 1 | Kamilar et al. 2010 | 0 | Webber and McGuire 2022 | 58205 | Myhrvold et al. 2015 | 885812 | Myhrvold et al. 2015 | 1600000 | Owen-Smith 1988 | 479 | Myhrvold et al. 2015 | 456 | Myhrvold et al. 2015 | 879 | Myhrvold et al. 2015 | 2069 | Myhrvold et al. 2015 |

Birth mass, weaning mass and female adult mass are reported in grams; female age at maturity, gestation length, lactation length and interbirth interval are reported in number of days.

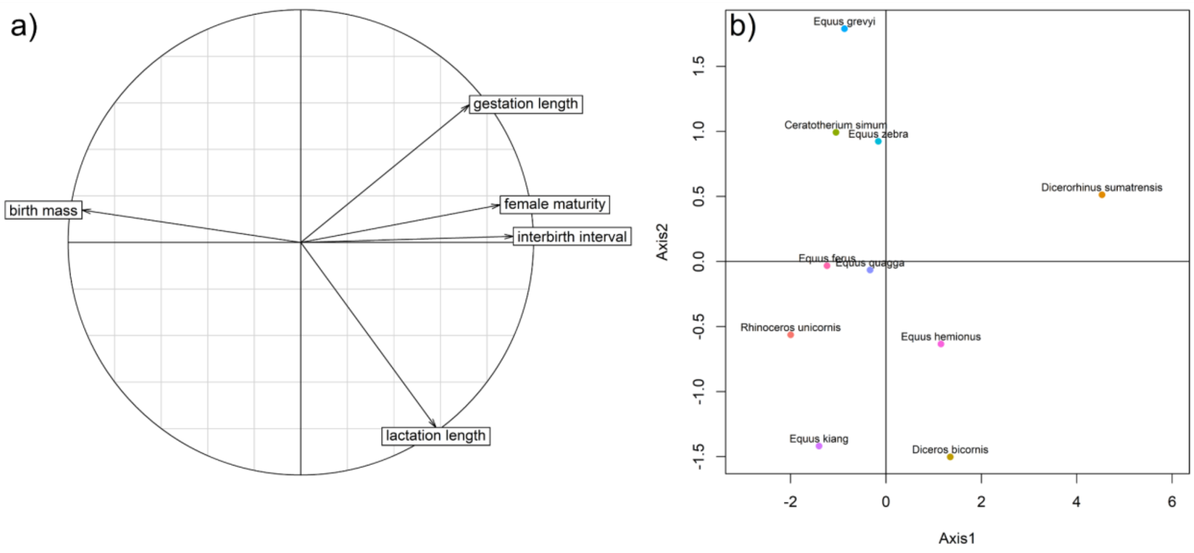


Figure S2.1: Principal Component Analysis (PCA) describing the pace of life of Perissodactyla species (*n* = 10 species) while correcting for differences in body mass: a) variables correlation circle, b) graph of individuals (here, individuals correspond to the 10 species of Perissodactyla). The variables included in the PCA correspond to the residuals of the regression of each life history trait (birth mass, gestation length, lactation length, interbirth interval and female age at maturity) according to the log-transformed female adult body mass. The first axis of the PCA explained 66% of the variance, the second axis explained 21% of the total variance.

Table S2.2: Variable loadings on the first and second axes of the principal component analysis describing the pace of life of Perissodactyla species (*n* = 10 species).

| **Variable** | **Loading on axis 1** | **Loading on axis 2** |
|---|---|---|
| Birth mass | -0.94 | 0.14 |
| Gestation length | 0.72 | 0.59 |
| Lactation length | 0.58 | -0.80 |
| Interbirth interval | 0.91 | 0.03 |
| Female age at maturity | 0.86 | 0.16 |

Supporting information 3: Determination of the predation exposure index.

***Determination of the predation risk***

The Jacob's and Ivelev's selectivity indices of a given predator species for a given prey species provide a quantitative estimate of the importance of a given prey species in the diet of a given predator species. They both vary between 1 (maximum preference) and -1 (maximum avoidance, Jacobs 1974). However, these indices can be unreliable as they vary spatially and temporally depending on the prey species available, they can easily be biased by data collection protocols (e.g. they tend to under-estimate rare and small preys due to infrequent or time-limited data collection sessions), and they were not available for all species and all sites considered in the present analyses at the time of the study. We thus used a combination of these selectivity indices as well as reports in the literature (with a special emphasis on juvenile sensitivity to the predator species considered) to attribute a qualitative level of predation risk specific to the combination of the Perissodactyla (prey species) and the predator species considered: 0 (no predation risk), 1 (low predation risk), 2 (medium predation risk), and 3 (high predation risk).

We considered the predation risk was absent if reported so in the studies or if no mention was found of the prey species in the diet of the predator species. We considered the predation risk was low if at least one study reported scattered evidence of predation attempts on the prey species or, in the absence of concrete evidence, if the prey species fell in the prey size range of the predator species and the absence of the prey species from the diet of the predator species was not explicitly stated. We considered the predation risk was medium if there was clear evidence of predation attempts on the prey species by the predator species in the literature. Finally, we considered the predation risk was high if several studies consistently reported the presence of the prey species in the diet of the predator species or, in the absence of multiple studies providing evidence, if the one study reported a high prevalence of the prey species in the diet of the predator species.

Despite being found in similar ecosystems as some of the Asian Perissodactyla retained in the analyses, the brown bear (*Ursus arctos*) was not described as predating on them (St-Louis and Côté 2009, Niedzialkowska et al. 2019). Although the golden jackal (*Canis aureus*) was mentioned as a potential predator of the onager (Shah and Qureshi 2007), we found no evidence of the presence of this Perissodactyla species in its diet in the literature and thus excluded this predator species.

Table S3.1: Variables used to determine the predation risk index for each species of Perissodactyla included in this study ($n$ = 10).

| **Perissodactyla species scientific name (reference)** | **Predator species scientific name (reference)** | **Jacob's index** | **Ivlev's index** | **Presence of Perissodactyla species in Predator species diet** | **Predation risk qualitative index** | **Citation** | **Reference (selectivity index)** | **Reference (presence in diet)** |
|---|---|---|---|---|---|---|---|---|
| *Ceratotherium simum* | *Acinonyx jubatus* | -1 | NA | 0 | 0 | not mentioned | Hayward et al. 2006b | / |
| *Diceros bicornis* | *Acinonyx jubatus* | -1 | NA | 0 | 0 | not mentioned | Hayward et al. 2006b | / |
| *Equus grevyi* | *Acinonyx jubatus* | NA | NA | 1 | 2 | *Nonetheless, cheetahs are known to take foals, and crocodiles and lions are known to take adults (Rowen and Ginsberg 1992).* | / | Moehlman 2002 (from other sources), https://animaldiversity.org/ |
| *Equus quagga* | *Acinonyx jubatus* | -0.69 | NA | 1 | 2 | *Cheetah diet was dominated by juvenile and neonate prey (67%) particularly of larger prey species like plains zebra, greater kudu, blue wildebeest, and waterbuck (Fig. 5b).* | Hayward et al. 2006b | Annear et al. 2023 |
| *Equus zebra* | *Acinonyx jubatus* | -1 | NA | 1 | 1 | / | Hayward et al. 2006b | https://animaldiversity.org/ |
| *Equus ferus* | *Canis lupus* | NA | 0.95 | 1 | 3 | (1) In Hustai National Park, Mongolia, the average survival rate of the released Przewalski's horses is 56.4%, and 51.95% of foals death was caused by wolf predation, 18.9% by inborn diseases and abortion, 8.8% by injury, 5% by insufficient milk intake, and 12.6% by unknown reasons. (2) 22.1% of mortalities were the result of attacks by wolves (3) Przewalski horses composed, 1% of wolf diets. | Lyngdoh et al. 2020 | (1) Chen et al. 2008, (2) Dorj and Namkhai 2013, (3) van Duyne et al. 2009 |
| *Equus hemionus* | *Canis lupus* | NA | NA | 1 | 3 | (1) Parts of hair or teeth of khulans and black-tailed gazelles (Gazella subgutturosa), the two large herbivores of Gobi B, occurred in 14 samples (40%). Direct observations on wolves' predatory behaviour on khulans were made on previous occasions (Feh et al., 1994). Furthermore, both during the observations in the winters of 1995 and 1996, wolf packs of four–five individuals pursued the regularly watched ''180 herd'' on several days, but without a successful kill. All authors having watched Mongolian khulans mention wolves as predators (Andrews, 1932, Bannikov, 1958, Zhirnov and Ilinski, 1986, Dulamtseren et al. 1989). Other predators were never observed or heard of attacking khulans. (2) Predation by the Indian grey wolf, which is the only predator on the wild ass, is very low, by 1983 there were only 65 wolves found in the area (Sinha, 1983) (3) As all animals observed were apparently in good body condition, predation by wolves provide the most likely explanation for the scarcity of young animals of the year. Wolf tracks along Kulan trails were seen daily. Furthermore, one direct observation of Wolf predation was made on August 6th 1993, from our hide. (4) Thus, there is no obvious mutual influence between mammals and the kulan, except for potential interactions with wolves. They are the only predators in the desert and steppe ecosystems of Central Asia that hunt the kulan. However, the quantitative predation of wolves on the kulan has not been studied, and it is believed that wolves mainly hunt foals, old or sick animals. Nevertheless, it can be assumed that the kulan will provide an additional food base for wolves. | / | (1) Feh et al. 2001, (2) Smielowski and Raval 1988, (3) Feh et al. 1994, (4) Abdibeka et al. 2025 |

| Perissodactyla species scientific name (reference) | Predator species scientific name (reference) | Jacob's index | Ivlev's index | Presence of Perissodactyla species in Predator species diet | Predation risk qualitative index | Citation | Reference (selectivity index) | Reference (presence in diet) |
|---|---|---|---|---|---|---|---|---|
| *Equus kiang* | *Canis lupus* | 0.91 | -0.13 | 1 | 1 | *Among potential predators, Tibetan wolf (Canis lupus chanco) and snow leopard (Uncia uncia) may occasionally prey on young and old individuals, but overall predation is unlikely to be an important limiting factor in populations of E. kiang (J. Van Gruisen, pers. Comm., Schaller 1998). Remains of E. kiang were absent in scats of C. lupus (n = 384), U. uncia (n 5 193), and Ursus arctos (Tibetan brown bear, n = 48) from Qinghai and Tibet (Schaller 1998). Observations from Mongolia report E. hemionus in scats of C. lupus (Feh et al. 2001).* | Shrotriya et al. 2022, Lyngdoh et al. 2020 | St-Louis and Côté 2009 |
| *Ceratotherium simum* | *Crocodylus niloticus* | NA | *NA* | 1 | 1 | *In the Complex the only predators capable of killing rhino are the lion Panthera leo, hyaena Crocuta crocuta and crocodile Crocodilus niloticus. There have been no records of lion or crocodile predation on black rhino although both species have been recorded killing square lipped rhino in the Complex.* | / | Hitchins and Anderson 1983 |
| *Diceros bicornis* | *Crocodylus niloticus* | NA | *NA* | 1 | 1 | *In the Complex the only predators capable of killing rhino are the lion Panthera leo, hyaena Crocuta crocuta and crocodile Crocodilus niloticus. There have been no records of lion or crocodile predation on black rhino although both species have been recorded killing square lipped rhino in the Complex.* | / | Hitchins and Anderson 1983 |
| *Equus grevyi* | *Crocodylus niloticus* | NA | *NA* | 1 | 1 | *Nonetheless, cheetahs are known to take foals, and crocodiles and lions are known to take adults (Rowen and Ginsberg 1992).* | / | Moehlman 2002 (from other sources) |
| *Equus quagga* | *Crocodylus niloticus* | NA | *NA* | 1 | 1 | *Crocodilians are exclusively carnivorous, and occupy the role of top predator in many tropical food webs (Mazzotti and Brandt, 1994, Lang, 2002) with the prey of adults including animals as large as wallabies, cattle, water buffalo, zebra, wildebeest, deer, and giraffe (Pye, 1976, Webb and Manolis, 1998, Shoop and Ruckdeschel, 1990, Shield, 1994, Alderton, 1998, Doody et al., 2007).* | / | Somaweera et al. 2013 (from other sources) |
| *Equus zebra* | *Crocodylus niloticus* | NA | *NA* | 0 | 0 | not mentioned | / | / |
| *Ceratotherium simum* | *Crocuta crocuta* | NA | NA | 0 | 1 | *(1) In the Complex the only predators capable of killing rhino are the lion Panthera leo, hyaena Crocuta crocuta and crocodile Crocodilus niloticus. There have been no records of lion or crocodile predation on black rhino although both species have been recorded killing square lipped rhino in the Complex. Although no actual attacks of hyaena on rhino calves have been witnessed, there is strong circumstantial evidence to suggest that this occurs frequently in Hluhluwe and to a lesser extent in the Corridor. (2) Neither at Umfolozi nor Hluhluwe did white rhino calves show torn ears to indicate attacks by hyenas.* | / | (1) Hitchins and Anderson 1983, (2) Owen-Smith 1988 |
| *Diceros bicornis* | *Crocuta crocuta* | NA | NA | 1 | 1 | (1) In the Complex the only predators capable of killing rhino are the lion Panthera leo, hyaena Crocuta crocuta and crocodile Crocodilus niloticus. There have been no records of lion or crocodile predation on black rhino although both species have been recorded killing square lipped rhino in the Complex. Although no actual attacks of hyaena on rhino calves have been witnessed, there is strong circumstantial evidence to suggest that this occurs frequently in Hluhluwe and to a lesser extent in the Corridor. (2) Rhinoceros appear to be vulnerable to predation by the spotted hyaena Crocuta crocuta Erxleben up to the age of about four months. Three attempts by hyaenas to pull down such animals were observed, all unsuccessful. (3) Black rhinos rarely feature in lion kill records, though there is an instance of a yearling killed and eaten by lions in the Serengeti. Hyena predation | / | (1) Hitchins and Anderson 1983, (2) Goddard 1967, (3) Owen-Smith 1988 |

| Perissodactyla species scientific name (reference) | Predator species scientific name (reference) | Jacob's index | Ivlev's index | Presence of Perissodactyla species in Predator species diet | Predation risk qualitative index | Citation | Reference (selectivity index) | Reference (presence in diet) |
|---|---|---|---|---|---|---|---|---|
| | | | | | | seems to be largely responsible for the poor survival of black rhino calves at Hluhluwe, and hyenas may be important predators on calves under 4 months old in East Africa. | | |
| *Equus grevyi* | *Crocuta crocuta* | 0.37 | NA | 1 | 2 | *On the LBL, endangered Grevy's zebra are favoured prey by both lions (Panthera leo) and spotted hyenas (Crocuta crocuta, hereafter referred to as hyenas), based on carcasses, scat and post hoc analyses of feeding sites identified by GPS telemetry data (Mwololo, 2011, Pratt, 2014, Dheer, 2016, Dupuis-Desormeaux et al., 2015).* | Davidson et al. 2019 | Davidson et al. 2019 |
| *Equus quagga* | *Crocuta crocuta* | -0.44, -0.24 | NA | 1 | 2 | *Lions are probably the most capable in handling equids and appear to prefer adult males, while hyenas and wild dogs appear to prefer adult female mountain zebra and common zebra (Berger 1983b and citings therein).* | Hayward 2006, Davidson et al. 2019 | Moehlman 2002 (from other sources) |
| *Equus zebra* | *Crocuta crocuta* | NA | NA | 1 | 2 | *Lions are probably the most capable in handling equids and appear to prefer adult males, while hyenas and wild dogs appear to prefer adult female mountain zebra and common zebra (Berger 1983b and citings therein).* | / | Moehlman 2002 (from other sources) |
| *Ceratotherium simum* | *Lycaon pictus* | NA | NA | 0 | 0 | not mentioned | / | / |
| *Diceros bicornis* | *Lycaon pictus* | NA | NA | 0 | 0 | not mentioned | / | / |
| *Equus grevyi* | *Lycaon pictus* | NA | NA | 1 | 1 | not mentioned | / | https://animaldiversity.org/ |
| *Equus quagga* | *Lycaon pictus* | -0.88 | NA | 1 | 1 | *(1) Lions are probably the most capable in handling equids and appear to prefer adult males, while hyenas and wild dogs appear to prefer adult female mountain zebra and common zebra (Berger 1983b and citings therein). (2) In all other studies small prey (< 50 kg) (impala, gazelle, or duiker) comprised the most important component of the diet (Fuller & Kat 1990: table 6). The only other studies that indicate substantial use of large prey are those of Pienaar (1969) which is discussed in detail below, and Malcolm & Van Lawick (1975) who describe specialization on zebra (Equus burchelli, 220–240 kg) by a single pack in the Serengeti.* | Hayward et al. 2006c | (1) Moehlman 2002 (from other sources), (2) van Dyk and Slotow 2003 (from other sources) |
| *Equus zebra* | *Lycaon pictus* | NA | NA | 1 | 1 | *Lions are probably the most capable in handling equids and appear to prefer adult males, while hyenas and wild dogs appear to prefer adult female mountain zebra and common zebra (Berger 1983b and citings therein).* | / | Moehlman 2002 (from other sources) |
| *Ceratotherium simum* | *Panthera leo* | -1 | NA | 1 | 1 | (1) In the Complex the only predators capable of killing rhino are the lion Panthera leo, hyaena Crocuta crocuta and crocodile Crocodilus niloticus. There have been no records of lion or crocodile predation on black rhino although both species have been recorded killing square lipped rhino in the Complex. Although no actual attacks of hyaena on rhino calves have been witnessed, there is strong circumstantial evidence to suggest that this occurs frequently in Hluhluwe and to a lesser extent in the Corridor (2) No instances of predation by lions or other carnivores were recorded during the study period (though two cases of white rhinos killed by lions have subsequently been noted: HITCHIN (pers. com.) and PIENAAR (1970) records an unsuccessful attack by lions on an adult bull in the Kruger Park). I watched lions in the close proximity of white rhinos on three | Hayward and Kerley 2005 | (1) Hitchins and Anderson 1983, (2) Owen-Smith 1975, (3) Owen-Smith 1988 |

| Perissodactyla species scientific name (reference) | Predator species scientific name (reference) | Jacob's index | Ivlev's index | Presence of Perissodactyla species in Predator species diet | Predation risk qualitative index | Citation | Reference (selectivity index) | Reference (presence in diet) |
|---|---|---|---|---|---|---|---|---|
| | | | | | | occasions, but the white rhinos seemed unconcerned and the lions disinterested. (3) A white rhino male was killed by lions in Umfolozi shortly after my departure (P. M. Hitchins, personal communication). From the Kruger Park in South Africa there are records of a white rhino calf killed by lions, and of a bull attacked and mauled so badly by lions that he had to be destroyed (Pienaar, 1970). | | |
| *Diceros bicornis* | *Panthera leo* | -1 | NA | 1 | 1 | (1) In the Complex the only predators capable of killing rhino are the lion Panthera leo, hyaena Crocuta crocuta and crocodile Crocodilus niloticus. There have been no records of lion or crocodile predation on black rhino although both species have been recorded killing square lipped rhino in the Complex. Although no actual attacks of hyaena on rhino calves have been witnessed, there is strong circumstantial evidence to suggest that this occurs frequently in Hluhluwe and to a lesser extent in the Corridor (2) The rhinoceros is vulnerable to predation by lion, even when adult (Ritchie, 1963). Proven cases are, however, rare. | Hayward and Kerley 2005 | (1) Hitchins and Anderson 1983, (2) Goddard 1967 |
| *Equus grevyi* | *Panthera leo* | 0.45 | NA | 1 | 3 | (1) On the LBL, endangered Grevy's zebra are favoured prey by both lions (Panthera leo) and spotted hyenas (Crocuta crocuta, hereafter referred to as hyenas), based on carcasses, scat and post hoc analyses of feeding sites identified by GPS telemetry data (Mwololo, 2011, Pratt, 2014, Dheer, 2016, Dupuis-Desormeaux et al., 2015). (2) Nonetheless, cheetahs are known to take foals, and crocodiles and lions are known to take adults (Rowen and Ginsberg 1992). | Davidson et al. 2019 | (1) Davidson et al. 2019, (2) Moehlman 2002 |
| *Equus quagga* | *Panthera leo* | 0.16, -0.20 | NA | 1 | 3 | *Lions are probably the most capable in handling equids and appear to prefer adult males, while hyenas and wild dogs appear to prefer adult female mountain zebra and common zebra (Berger 1983b and citings therein).* | Hayward and Kerley 2005, Davidson et al. 2019 | Moehlman 2002 (from other sources) |
| *Equus zebra* | *Panthera leo* | NA | NA | 1 | 3 | *Lions are probably the most capable in handling equids and appear to prefer adult males, while hyenas and wild dogs appear to prefer adult female mountain zebra and common zebra (Berger 1983b and citings therein).* | / | Moehlman 2002 (from other sources) |
| *Ceratotherium simum* | *Panthera pardus* | -1 | NA | 0 | 0 | not mentioned | Hayward et al. 2006a | / |
| *Diceros bicornis* | *Panthera pardus* | -1 | NA | 0 | 0 | not mentioned | Hayward et al. 2006a | / |
| *Equus grevyi* | *Panthera pardus* | NA | NA | 1 | 1 | not mentioned | / | https://animaldiversity.org/ |
| *Equus quagga* | *Panthera pardus* | -0.80 | NA | 1 | 1 | *Foals are preyed upon by lions, spotted hyenas, cheetahs and leopards.* | Hayward et al. 2006a | https://www.sanbi.org/ |
| *Equus zebra* | *Panthera pardus* | -1 | NA | 1 | 1 | not mentioned | Hayward et al. 2006a | https://animaldiversity.org/ |
| *Dicerorhinus sumatrensis* | *Panthera tigris* | NA | NA | 1 | 2 | *Sumatran rhinoceroses do not have many predators, except for humans (Homo sapiens) and Sumatran tigers (Panthera tigris sumatrae). Sumatran tigers do not actively hunt Sumatran rhinoceroses, but will opportunistically attack stranded calves.* | / | https://animaldiversity.org/ |

| **Perissodactyla species scientific name (reference)** | **Predator species scientific name (reference)** | **Jacob's index** | **Ivlev's index** | **Presence of Perissodactyla species in Predator species diet** | **Predation risk qualitative index** | **Citation** | **Reference (selectivity index)** | **Reference (presence in diet)** |
|---|---|---|---|---|---|---|---|---|
| *Equus hemionus* | *Panthera tigris* | NA | NA | 0 | 0 | *Tigers also hunt kulans, but a serious level of preying is not expected, since they will occupy generally different habitats (kulans – open areas, tigers - fairly dense plantings), encountering only at watering places.* | / | Abdibeka et al. 2025 |
| *Rhinoceros unicornis* | *Panthera tigris* | -1 | NA | 1 | 2 | (1) Causes of death included poaching, tiger predation on the calves and intraspecific fighting among the males. Predation by tigers on calves is the most likely form of death to go undetected. The fact that I recorded three such deaths implies that tiger predation is quite common (2) Amongst the 47 calf deaths recorded, majority (29.8%) were due to tiger predation (3) All calf mortality (n = 4) occurred during the first year of life when calves were prey for tigers (4) The predation of Rhino calf by tiger is a common phenomenon in Rajiv Gandhi Orang National Park as the Rhino calfs are found throughout the year (5) Commonly, tigers are known for their predation on herbivores other than rhinos, but in Kaziranga National Park, the Royal Bengal tiger preyed upon 203 rhino calves between 1985-2000. | Hayward et al. 2012 | (1) Laurie 1978, (2) Sudebi et al. 2017, (3) Dinerstein and Price 1991, (4) Hazarika and Saikia 2010, (5) Talukdar 2002 |
| *Equus ferus* | *Panthera uncia* | -0.31 | NA | 0 | 0 | not mentioned | Lyngdoh et al. 2014 | / |
| *Equus hemionus* | *Panthera uncia* | -1 | NA | 0 | 0 | not mentioned | Lyngdoh et al. 2014 | / |
| *Equus kiang* | *Panthera uncia* | -1 | NA | 1 | 1 | *Among potential predators, Tibetan wolf (Canis lupus chanco) and snow leopard (Uncia uncia) may occasionally prey on young and old individuals, but overall predation is unlikely to be an important limiting factor in populations of E. kiang (J. Van Gruisen, pers. Comm., Schaller 1998). Remains of E. kiang were absent in scats of C. lupus (n = 384), U. uncia (n 5 193), and Ursus arctos (Tibetan brown bear, n = 48) from Qinghai and Tibet (Schaller 1998). Observations from Mongolia report E. hemionus in scats of C. lupus (Feh et al. 2001).* | Shrotriya et al. 2022 | St-Louis and Côté 2009 |

***Determination of the predator abundance***

Predator densities can be estimated using different methods providing non-comparable estimates (Whittington and Sawaya 2015, Alexander et al. 2016, Morin et al. 2022), fluctuate through time (e.g. Trinkel 2013), and are not precisely quantified in some study sites (e.g. van Duyne et al. 2009). Therefore, we decided to quantify modern (i.e. between the 1900's and nowadays) predator densities in each study site semi-quantitatively and relatively to the species usual abundance level as follows: 0 (absent from the study site), 1 (low abundance), 2 (medium abundance), 3 (high abundance) (Table S3.2). To do so, we first extracted references of predator densities for each of our study sites. When no data was available for the study site of interest, we used data from surrounding areas. We used various sources from site- and predator-specific, non-systematic searches on Google Scholar such as original and review scientific papers as well as empirical observations from technical reports and theses, and as a last resort, list of species provided by the parks for tourism purposes. When available, we also used mentions of predator population estimates directly from the studies providing distributions of births in Perissodactyla. We reported four descriptors of predator abundance, sorted from the most to the least accurate: (1) density estimates, (2) population counts (converted into an estimate of number of individuals per 100 $km^2$ based on the surface of the study site), (3) empirical reports of abundance, (4) reports of presence or absence of the species in the study site. Using all the predator densities and the GPS coordinates extracted by Santini et al. (2018), we calculated a site-specific mean predator density. Snow leopard (*Panthera uncia*) and Nile crocodile (*Crocodylus niloticus*) were not included in this review article, so we extracted densities from various sources and used a similar approach as described above (Table S3.3). We then calculated the 0.33 and 0.66 percentiles of the distribution of densities for each species of predator. Finally, we compared the densities we retrieved for our study sites of interest using descriptors (1) and (2) (i.e. density estimates and population counts converted into an estimate of number of individuals per 100 $km^2$) with the percentiles of the density distributions and attributed the semi-quantitative index as follows:

- density <0.33 percentile: low abundance
- density <0.66 and >0.33 percentiles: medium abundance
- density >0.66 percentile: high abundance

When only descriptors (3) and (4) were available (i.e. empirical reports of abundance and reports of presence or absence of the species), we attributed a medium abundance, unless specified otherwise in the reference ($n = 4$, e.g. "*Records for wild dogs* Lycaon pictus *are rare and they have been described as being present only intermittently*", Oates and Rees 2013). If the predator species was reported as rare, we attributed a low abundance, and if the predator was reported as numerous, we attributed a high abundance.

Table S3.2: Site-specific predator abundance index for each species of predator included in this study (*n* = 9).

| Predator species scientific name (reference) | Study site name (reference) | Quote | Density (number of individuals /100 km$^2$) | Study site (if different from reference) | Reference | Predator density qualitative index |
|---|---|---|---|---|---|---|
| *Acinonyx jubatus* | Addo Elephant National Park, South Africa | [species not reported] | 0.00 | | https://www.sanparks.org/ | 0 |
| *Acinonyx jubatus* | Etosha National Park and Daan Viljoen Game Reserve and Namib Desert Park and Naukluft Mountain Zebra Park, Namibia | (1) 0.20 Individual per 100 km2 (estimated). (2) We identified 30 [...] individual cheetahs | 0.20 | | (1) Weise et al. 2017, (2) Keja et al. 2025 | 1 |
| *Acinonyx jubatus* | Hluhluwe Corridor Umfolozi Game Reserve Complex, South Africa | (1) However, the extirpation of the other members of the large carnivore guild in the first half of the 20th century has meant that [...] cheetahs (Acinonyx jubatus) have all had to be reintroduced (Cromsigt et al. 2017; Trinkel et al. 2008). The estimated numbers of these latter three species in 2018 were [...] eight [cheetah] (Roberts 2022). (2) Cheetah presence documented | 0.83 | | (1) Roberts et al. 2023, (2) Davis et al. 2024 | 2 |
| *Acinonyx jubatus* | Hwange National Park, Zimbabwe | *Mean population densities of resident cheetah populations were highest in conservancies and lowest in parastatal wildlife estates (Table 2), with a maximum density of 2.50 ind/100 km2 in the Malilangwe conservancy (Fig. 1) and a minimum density of 0.15 ind/100 km2 in Hwange National Park.* | 0.15-2.50 | | van der Meer 2018 | 2 |
| *Acinonyx jubatus* | Ithala Game Reserve, South Africa | *IGR has been virtually predator free since its creation in 1972.* | 0.00 | | O'Kane and Macdonald 2016 | 0 |
| *Acinonyx jubatus* | Kruger National Park, South Africa | *A total of 412 (329–495; SE 41.95) cheetahs [...] occur in the Kruger National Park.* | 2.11 | | Marnewick et al. 2014 | 3 |
| *Acinonyx jubatus* | Lake Mutirikwi Recreational Park, Zimbabwe | *There are no large predators in the Park apart from the indigenous side-striped jackal (Canis adustus). Pythons (Python sebae), and crocodiles (Crocodylus niloticus) which are numerous in the lake, may remove a few animals although very few such records exist.* | 0.00 | | Condy 1973 | 0 |
| *Acinonyx jubatus* | Lewa Wildlife Conservancy, Kenya | Presence reported on camera trap images | NA | | Sargent 2016 | 2 |
| *Acinonyx jubatus* | Masai Mara National Reserve, Kenya | *We estimate adult cheetah density to be between 1.28 ± 0.315 and 1.34 ± 0.337 individuals/100km2 across four candidate models specified in our analysis.* | 1.28-1.34 | | Broekhuis and Gopalaswamy 2016 | 2 |
| *Acinonyx jubatus* | Mountain Zebra National Park, South Africa | *Cheetah Acinonyx jubatus have also been reintroduced MZNP, as well as into the areas of three additional sub-populations. [...] Cheetah are thought to be limiting the population growth in at least one private property where 13 partially consumed CMZ carcasses were found over an 18 month period. The cause of death could not be confirmed for these individuals, but cheetah numbers exceeded 21 individuals during that time.* | 7.39 | | Hrabar and Kerley 2015 | 3 |
| *Acinonyx jubatus* | Ngorongoro Crater, Tanzania | *Cheetahs Acinonyx jubatus have been described as being present intermittently in the crater (Homewood & Rodgers 1991) and it has been suggested that they were absent prior to c. 1988 (Martin 1998). However, Barns recorded their presence in 1921 (Barns 1921) and GHR Hurst described cheetahs as 'common' (Joelson 1928). Cheetahs were described by Laurenson (1991) as 'virtually absent', although they were recorded in 2004–06* | NA | | Oates and Rees 2013 | 1 |

| Predator species scientific name (reference) | Study site name (reference) | Quote | Density (number of individuals /100 km$^2$) | Study site (if different from reference) | Reference | Predator density qualitative index |
|---|---|---|---|---|---|---|
| *Acinonyx jubatus* | North, Namibia | 0.19 ind/100km2 in Zambezian Baikiaea Woodlands, 0.70 ind/100km2 in Kalahari Xeric Savanna and Gariep Karoo, 0.20 ind/100km2 in Namibian Savanna Woodland, Namib Desert and Gariep Karoo. | 0.19-0.70 | | Weise et al. 2017 (from other sources) | 1 |
| *Acinonyx jubatus* | Pilanesberg National Park, South Africa | *Other large predators include [...] cheetahs (about 20)* | 3.50 | | van Dyk and Slotow 2003 | 3 |
| *Acinonyx jubatus* | Save Valley Conservancy and Imire Game Ranch, Zimbabwe | *Populations of cheetah [...] already existed in the area when SVC was formed [...] (Lindsey et al., 2009b)* | NA | | Williams 2011 | 2 |
| *Acinonyx jubatus* | Serengeti, Tanzania | between 0.02 and 0.12 individuals per km2 [From Figure 3] | 2.00-12.00 | | Durant et al. 2011 | 3 |
| *Acinonyx jubatus* | Sinamatella Intensive Protection Zone, Zimbabwe | [same reports as Hwange National Park, Zimbabwe considered for this study site] | 0.15-2.50 | | / | 2 |
| *Acinonyx jubatus* | Tsavo East National Park, Kenya | *cheetah (154 ± 74)* | 1.12 | Tsavo National Parks | Henschel et al. 2020 | 2 |
| *Acinonyx jubatus* | Umfolozi Game Reserve, South Africa | [same reports as Hluhluwe Corridor Umfolozi Game Reserve Complex, South Africa considered for this study site] | 0.83 | | / | 2 |
| *Acinonyx jubatus* | Zambia | *(1) Historically the cheetah was recorded as a widespread but rare to uncommon species in Zambia (Myers 1975, Ansell 1978). Cheetahs were resident in most protected areas, the Lower Zambezi complex being the only exception, where cheetahs were recorded as either absent (Skinner & Smithers, 1990) or as vagrants (Ansell 1978, Nowell & Jackson 1996, Skinner & Chimimba, 2005; Fig. 1). (2) A small but stable population of cheetah (15–20 known individuals) is present.* | 0.52 | (2) Western Zambia's Greater Liuwa Ecosystem | (1) Purchase 2007, (2) Martens et al. 2025 | 1 |
| *Acinonyx jubatus* | Ziwa Rhino Sanctuary, Uganda | [species not reported] | 0.00 | | https://ziwarhinoandwildliferanch.com/about-ziwa/wildlife-on-the-ranch/ | 0 |
| *Canis lupus* | Chang Tang Reserve, China | *With the alpine steppe now essentially usurped by pastoralists, the northern part of the reserve represents the last real refuge for wildlife and especially for the wild yak, wolf, and bear.* | NA | | Miller and Schaller 1996 | 2 |
| *Canis lupus* | Chitwan National Park, Nepal | [species not reported] | 0.00 | | / | 0 |
| *Canis lupus* | Gobi A National Park, Mongolia | *We collected wolf (Canis lupus) droppings from 35 different sites on random itineraries throughout Gobi B* | NA | | Feh et al. 2001 | 2 |
| *Canis lupus* | Hustai National Park, Mongolia | *The HNP staff and the herdsmen around the park claim that there are about 50 wolves living in the park (P. Enkhkhuyag, personal communication). We estimated the population at 20–50 individuals (C. van Duyne and E. Ras, Wageningen University, unpublished data), based on calculations that included wolf energy requirements (Nagy 1987), interview data, and scat analysis.* | 6.92 | | van Duyne et al. 2009 | 3 |

| Predator species scientific name (reference) | Study site name (reference) | Quote | Density (number of individuals /100 km$^2$) | Study site (if different from reference) | Reference | Predator density qualitative index |
|---|---|---|---|---|---|---|
| *Canis lupus* | Takhin Tal, Mongolia | *In 2002/03, 60 wolves were killed in the Bugat, Tonkhil and Altai-Khovd part of the study area (15,912 km2), in 2003/04, 46 were killed in the Bugat and Tonkhil part of the study area (7,074 km2) and in 2004/05, 78 were killed in the Bugat, Tonkhil, Altai-Khovd and Uench part of the study area (18,670 km2). This corresponds to a harvest rate of roughly 1 wolf/265 km2 in 2002/03, 1 wolf/120 km2 in 2003/04 and 1 wolf/310 km2 in 2004/05. However, hunting pressure was unevenly distributed and was particularly high in the north and northeastern part of the park. During the active monitoring period of F1 in 2003/04 and 2004/05, 80 wolves were killed within the area of her multi-annual home range (1 wolf/80 km2) and 34 wolves (1 wolf/38 km2) within her smaller 'resident' range (see Fig. 3).* | NA | Great Gobi B SPA | Kaczensky et al. 2008 | 2 |
| *Canis lupus* | Xinjiang, China | *Eight wolves that represented four distinct populations from lowland (Xinjiang and Inner Mongolia) and highland (Tibet and Qinghai) (Fig. 1 and S2; Table S1) were selected for genome sequencing* | NA | | Zhang et al. 2014 | 2 |
| *Crocodylus niloticus* | Addo Elephant National Park, South Africa | [species not reported] | 0.00 | | https://www.sanparks.org/ | 0 |
| *Crocodylus niloticus* | Etosha National Park and Daan Viljoen Game Reserve and Namib Desert Park and Naukluft Mountain Zebra Park, Namibia | [species not reported] | 0.00 | | / | 0 |
| *Crocodylus niloticus* | Hluhluwe Corridor Umfolozi Game Reserve Complex, South Africa | *Despite being one of the oldest protected areas in Africa, formally established in 1895 and once the exclusive royal hunting ground of King Shaka, we are not aware of a Nile crocodile aerial survey ever conducted in HIP. In 1982, Pooley recorded 144 nests (Hartley 1990). Even though the population at the time was not known, assuming a 50% nest effort (likely to be considerably lower) and an even sex ratio (1:1), which is known for nearby Lake St Lucia (Warner et al. 2016b), then we could assume 288 females and an adult population of 576 crocodiles. Hartley (1990) continued with nest surveys from 1984-1987 and found a mean number of 108.3 ± 20.4 nests along the White and Black uMfolozi Rivers, a combined distance of 124.5 km.* | 60.00 | | Combrink et al. 2019 | 3 |
| *Crocodylus niloticus* | Hwange National Park, Zimbabwe | [species reported] | NA | | Utete 2021 | 2 |
| *Crocodylus niloticus* | Ithala Game Reserve, South Africa | [species reported] | NA | | https://www.nature-reserve.co.za/ithala-game-reserve-birding-game-viewing.html | 2 |
| *Crocodylus niloticus* | Kruger National Park, South Africa | *(1) Even so, corrected spotlight- and helicopter-based estimates were comparable and the number of crocodiles in the focal study area declined significantly from 780 (95% CI: 637–1222) to between 460 (spotlight estimate, 95% CI 375–665) and 505 (aerial estimate, 95% CI: 559–1746) during the period of crocodile deaths. (2) KNP, which also occurs in Mpumalanga Province, continues to be the most important protected area for Nile crocodiles in South Africa. In 2017, 3326 individuals were counted from a helicopter, the third highest count since helicopter surveys commenced in 1982. The severe drought in 2015 and 2016 created optimal counting conditions and numerous crocodiles were counted in dams and pans as streams and seasonal rivers dried up completely (SANParks survey data). During the height of the* | 17.07 | | (1) Ferreira and Pienaar 2011, (2) Combrink et al. 2019 | 2 |

| Predator species scientific name (reference) | Study site name (reference) | Quote | Density (number of individuals /100 km$^2$) | Study site (if different from reference) | Reference | Predator density qualitative index |
|---|---|---|---|---|---|---|
| | | *drought a number of crocodiles were isolated and died along the Shingwedzi River (D. Pienaar, pers. comm.).* | | | | |
| *Crocodylus niloticus* | Lake Mutirikwi Recreational Park, Zimbabwe | *There are no large predators in the Park apart from the indigenous side-striped jackal (Canis adustus). Pythons (Python sebae), and crocodiles (Crocodylus niloticus) which are numerous in the lake, may remove a few animals although very few such records exist.* | NA | | Condy 1973 | 3 |
| *Crocodylus niloticus* | Lewa Wildlife Conservancy, Kenya | [species not reported] | 0.00 | | / | 0 |
| *Crocodylus niloticus* | Masai Mara National Reserve, Kenya | *For instance, the crocodile often considered a very dangerous animal and feared by many people, accounted for few of the conflict incidences in Tsavo (0.6%, n = 199) and Mara (0.1%, n = 14). The few cases of crocodile conflicts in these two regions do not necessarily reflect the national threat posed to humans and livestock by crocodiles and may reflect the fact that crocodiles inhabit sections of large rivers within protected areas with low human and livestock populations.* | NA | | Mukeka et al. 2018 | 2 |
| *Crocodylus niloticus* | Mountain Zebra National Park, South Africa | [species not reported] | 0.00 | | https://www.sanparks.org/ | 0 |
| *Crocodylus niloticus* | Ngorongoro Crater, Tanzania | [species not reported] | 0.00 | | / | 0 |
| *Crocodylus niloticus* | North, Namibia | *The Nile crocodile is seen as an integral component of the aquatic ecosystems of northeastern (Caprivi region) and northwestern (Kunene River) Namibia. [...] Mean total (ie uncorrected) counts were: Kwando River 37 ± 16, Mamili National Park 38 ± 7, Okavango River 29 ± 10, Linyanti/Chobe Rivers 60 ± 30 and Zambezi River 45 ± 18. Concealment and spotlight-to-aerial count correction factors were applied and resulted in the following population estimates: Kwando River 1379 ± 605, Mamili National Park 1328 ± 251, Okavango River 912 ± 318, Linyanti/Chobe Rivers 578 ± 286 and Zambezi River 117 ± 47 crocodiles.* | NA | Caprivi region and Kunene River | Combrink et al. 2019 | 2 |
| *Crocodylus niloticus* | Pilanesberg National Park, South Africa | Mean count in 2012: 19 +/- 6, in 2013: 33 +/- 3 | 4.53 | | Power and Verburgt 2014 | 1 |
| *Crocodylus niloticus* | Save Valley Conservancy and Imire Game Ranch, Zimbabwe | (1) Aerial surveys using a two-seater fixed-wing aeroplane (pilot plus one observer/recorder) were conducted to record Nile crocodiles in 2008-2011 in the Savé, Runde and Mwenezi Rivers (Zisadza-Gandiwa et al. 2013). A total of 307 ± 44.6 SD individuals was recorded over the four years, with the Runde River hosting most crocodiles (240 ± 41.2 SD), followed by the Mwenezi River (45.3 ± 1.9 SD) and Savé River (22 ± 3.9 SD). (2) Results showed that lions (Panthera leo), spotted hyenas (Crocuta crocuta), elephants (Loxodonta africana), and Nile crocodiles (Crocodylus niloticus) were the major animals involved in the conflict. | 9.19 | | (1) Combrink et al. 2019, (2) Makumbe et al. 2022 | 2 |
| *Crocodylus niloticus* | Serengeti, Tanzania | The predators considered were lions and crocodiles. | NA | | Ngana et al. 2014 | 2 |
| *Crocodylus niloticus* | Sinamatella Intensive Protection Zone, Zimbabwe | [same reports as Hwange National Park, Zimbabwe considered for this study site] | NA | | / | 2 |
| *Crocodylus niloticus* | Tsavo East National Park, Kenya | [species reported] | NA | | https://www.tsavonationalparkkenya.com | 2 |

| Predator species scientific name (reference) | Study site name (reference) | Quote | Density (number of individuals /100 km$^2$) | Study site (if different from reference) | Reference | Predator density qualitative index |
|---|---|---|---|---|---|---|
| *Crocodylus niloticus* | Umfolozi Game Reserve, South Africa | [same reports as Hluhluwe Corridor Umfolozi Game Reserve Complex, South Africa considered for this study site] | 60.00 | | / | 3 |
| *Crocodylus niloticus* | Zambia | *Wallace et al. (2013) conducted two spotlight surveys, in 2006 and 2009, to estimate population size, structure and trends for wild Nile crocodiles in the lower/middle Zambezi Valley, the stretch of the Zambezi River between Lake Kariba and Cahora Bassa Dam in Mozambique, covering a distance of approximately 270 km. This area is important for conservation as well as being a source of crocodile eggs for the ranching industry. A total of 1761 crocodiles were encountered during the spotlight survey in 2009, 51.5% juveniles, 29.7% sub-adults, 18.8% adults and 33.8% eyes-only, giving a total 2009 estimate (including correction factors) of 2257 crocodiles. For the purpose of comparison, the 2006 population estimate was corrected to 1984 crocodiles (using the 2009 riverbank measurements). The survey data suggest an increase in population density from 2.2 km-1 in 2006 to 2.5 km-1 in 2009. Crocodile density was greatest (5.4 km-1) in the areas of wildlife and habitat protection (Mana Pools) and lowest (0.9 km-1) in areas of increased human presence (Siavunga open area).* | NA | | Combrink et al. 2019 | 2 |
| *Crocodylus niloticus* | Ziwa Rhino Sanctuary, Uganda | [species not reported] | 0.00 | | https://ziwarhinoandwildliferanch.com/about-ziwa/wildlife-on-the-ranch/ | 0 |
| *Crocuta crocuta* | Addo Elephant National Park, South Africa | *(1) This is probably because of the low density of each (0.04 lions and 0.07 hyaenas km-2), reducing the likelihood of competitive or agonistic interactions. (2) eight spotted hyaenas (five males and three females) were reintroduced to the Main Camp section of the park in 2003 and 2004 (Hayward et al. 2007a, c)* | 7.00 | | (1) Hayward and Hayward 2006, (2) Hayward et al. 2009 | 1 |
| *Crocuta crocuta* | Etosha National Park and Daan Viljoen Game Reserve and Namib Desert Park and Naukluft Mountain Zebra Park, Namibia | *Based on this mathematical correlation, I estimated 203 +/- 79 hyenas, i.e., 2.7 +/- 1.1 hyenas/100 km2, in the central and eastern parts of Etosha. Applying this correlation to the western part of the park, it was possible to estimate 339 +/- 176 hyenas, corresponding to an overall density of 2.1 +/- 1.0 hyenas/100 km2, in the whole Etosha National Park.* | 1.00 | | Trinkel 2009 | 1 |
| *Crocuta crocuta* | Hluhluwe Corridor Umfolozi Game Reserve Complex, South Africa | *(1) Average hyaena density, at 0.357 individuals/km2, was relatively high compared to other southern African conservation areas, and range from 0 to 1.25 individuals/km2 across sampling stations. (2) We estimated an average of 18.29 ± 3.27 spotted hyaenas per 100 km2 between 2013 and 2018, with an annual estimated high of 20.83/100 km2 in 2014 and a low of 11.98/100 km2 in 2015.* | 18.29 | | (1) Graf et al. 2009, (2) Roberts et al. 2023 | 2 |
| *Crocuta crocuta* | Hwange National Park, Zimbabwe | *In the study area, the average hyaena density between 2009 and 2012 was 9.2 hyaenas/100 km2 [...] (Andrew J. Loveridge, pers. com).* | 9.20 | | Périquet et al. 2016 | 1 |
| *Crocuta crocuta* | Ithala Game Reserve, South Africa | *IGR has been virtually predator free since its creation in 1972.* | 0.00 | | O'Kane and Macdonald 2016 | 0 |
| *Crocuta crocuta* | Kruger National Park, South Africa | 2340 individuals in 1985, 2992 individuals in 1989, 3348 individuals in 2005/2006, 3667 individuals in 2008 | 15.84 | | Ferreira and Funston 2016 | 2 |

| Predator species scientific name (reference) | Study site name (reference) | Quote | Density (number of individuals /100 km$^2$) | Study site (if different from reference) | Reference | Predator density qualitative index |
|---|---|---|---|---|---|---|
| *Crocuta crocuta* | Lake Mutirikwi Recreational Park, Zimbabwe | *There are no large predators in the park apart from the indigenous side-striped jackal (Canis adustus). Pythons (Python sebae), and crocodiles (Crocodylus niloticus) which are numerous in the lake, may remove a few animals although very few such records exist.* | 0.00 | | Condy 1973 | 0 |
| *Crocuta crocuta* | Lewa Wildlife Conservancy, Kenya | Presence reported on camera trap images | NA | | Sargent 2016 | 2 |
| *Crocuta crocuta* | Masai Mara National Reserve, Kenya | *Apparent hyena density estimates were 1.3 times higher in the ranches (0.561 hyenas/km2) than in the reserve (0.404 hyenas/km2), in correspondence with the regional pattern of prey density. This distribution of hyenas is biased towards the reserve, if it is dependent on prey density. [...] Lion and hyena densities and prey biomass did not differ between June 1991 (5172.273 kg/km2) and June 2003 (5472 kg/km2) in the reserve [...].* | 40.40-56.10 | | Ogutu at al. 2005 | 3 |
| *Crocuta crocuta* | Mountain Zebra National Park, South Africa | [species not reported] | 0.00 | | https://www.sanparks.org/ | 0 |
| *Crocuta crocuta* | Ngorongoro Crater, Tanzania | *(1) The number of adult hyenas in the Crater in 1996 was 117, 2.5 times lower than the 298 adult hyenas (minimum estimate), or 3.3 times lower than the 385 adult hyenas (Lincoln index) estimated to inhabit the Crater in the late 1960s. (2) Data on the spotted hyena population size are rare. Kruuk (1972) estimated the total at 430 individuals, including 385 adults. By 2000, the population had apparently declined to 139 (Estes et al. 2006).* | 53.46-148.08 | | (1) Höner et al. 2005, (2) Oates and Rees 2013 | 3 |
| *Crocuta crocuta* | North, Namibia | *SPACECAP provided a summary of results (Table 1) and determined a density of 0.85 hyaenas/100 km2 for the TRV (SD = 0.14). Furthermore, SPACECAP also provided an output file describing pixel densities. These pixel densities were used to generate a pixelated map showing fine-scale variation in density (Figure 1). Each pixel represents a density of 0.25 km2 and ranges between 0.01/100 and >0.05/100 km2. The highest density of pixels is located in the southern lower lying canyons, furthest away from any anthropogenic features, such as property B (Figure 1).* | 0.85 | Tsauchab River Valley | Fouché et al. 2020 | 1 |
| *Crocuta crocuta* | Pilanesberg National Park, South Africa | *The reserve has lions but no spotted hyenas.* | 0.00 | | van Dyk and Slotow 2003 | 0 |
| *Crocuta crocuta* | Save Valley Conservancy and Imire Game Ranch, Zimbabwe | *Results showed that lions (Panthera leo), spotted hyenas (Crocuta crocuta), elephants (Loxodonta africana), and Nile crocodiles (Crocodylus niloticus) were the major animals involved in the conflict.* | NA | | Makumbe et al. 2022 | 2 |
| *Crocuta crocuta* | Serengeti, Tanzania | between 0.16 and 2.55 individuals per km2 [From Figure 3] | 16.00-255.00 | | Durant et al. 2011 | 3 |
| *Crocuta crocuta* | Sinamatella Intensive Protection Zone, Zimbabwe | [same reports as Hwange National Park, Zimbabwe considered for this study site] | 9.20 | | / | 1 |
| *Crocuta crocuta* | Tsavo East National Park, Kenya | *Spotted hyaenas were the most abundant, with an estimated population of 3,903 ± 514 (95% CI)* | 28.39 | Tsavo National Parks | Henschel et al. 2020 | 2 |
| *Crocuta crocuta* | Umfolozi Game Reserve, South Africa | [same reports as Hluhluwe Corridor Umfolozi Game Reserve Complex, South Africa considered for this study site] | 18.29 | | / | 2 |
| *Crocuta crocuta* | Zambia | *(1) We identified 663 individual hyenas from 11 clans and one group of individuals not assigned to a clan at the end of the study, across the GLE. (2) We estimated a minimum density of 0.13 hyaenas/km2 and* | 13.00-52.00 | (1) Western Zambia's | (1) Martens et al. 2025, (2) | 2 |

| Predator species scientific name (reference) | Study site name (reference) | Quote | Density (number of individuals /100 km$^2$) | Study site (if different from reference) | Reference | Predator density qualitative index |
|---|---|---|---|---|---|---|
| | | *maximum density of 0.52 hyaenas/km2. Density fluctuated with seasonal migrations of the main prey species, the blue wildebeest.* | | Greater Liuwa Ecosystem, (2) Liuwa Plain National Park | M'soka et al. 2016 | |
| *Crocuta crocuta* | Ziwa Rhino Sanctuary, Uganda | [species not reported] | 0.00 | | https://ziwarhinoandwildliferanch.com/about-ziwa/wildlife-on-the-ranch/ | 0 |
| *Lycaon pictus* | Addo Elephant National Park, South Africa | [species not reported] | 0.00 | | https://www.sanparks.org/ | 0 |
| *Lycaon pictus* | Etosha National Park and Daan Viljoen Game Reserve and Namib Desert Park and Naukluft Mountain Zebra Park, Namibia | *An attempted reintroduction of captive wild dogs into Etosha National Park failed when a pride of lions systematically hunted and killed the dogs over several weeks (L. Scheepers, personal communication).* | 0.00 | | Creel and Creel 1996 | 0 |
| *Lycaon pictus* | Hluhluwe Corridor Umfolozi Game Reserve Complex, South Africa | *(1) From 2006 to 2009 the Hluhluwe-iMfolozi wilddog population increased from 44 animals in six packs to 95 animals in eight packs. (2) However, the extirpation of the other members of the large carnivore guild in the first half of the 20th century has meant that African lions (Panthera leo), African wild dogs (Lycaon pictus), and cheetahs (Acinonyx jubatus) have all had to be reintroduced (Cromsigt et al. 2017; Trinkel et al. 2008). The estimated numbers of these latter three species in 2018 were [...] 36 [wild dog] [...] (Roberts 2022).* | 3.75-9.90 | | (1) Whittington-Jones et al. 2014, (2) Roberts et al. 2023 | 3 |
| *Lycaon pictus* | Hwange National Park, Zimbabwe | *(1) In 2004 it was estimated that African wild dog densities in and around Hwange National Park were 1.5 African wild dogs/100km2 which is around 225 individuals (Woodroffe et al. 2004). More recent information suggests that there are approximately 50 to 70 individuals left (Zimbabwe Parks and Wildlife Management Authority 2009, Blinston 2010). (2) 18 ind/1000 km2 [From Table 1]* | 0.34-1.80 | | (1) van der Meer 2011, (2) Fuller et al. 1992 | 1 |
| *Lycaon pictus* | Ithala Game Reserve, South Africa | *IGR has been virtually predator free since its creation in 1972.* | 0.00 | | O'Kane and Macdonald 2016 | 0 |
| *Lycaon pictus* | Kruger National Park, South Africa | (1) Overall 357 wild dogs were identified, which represents the minimum number present in the study area at the beginning of 1989. This gives a minimum density of 16.7 dogs/1000 km 2, or 1 dog/60 km 2. (2) A total of [...] 151 (144–157; SE 3.21) wild dogs occur in the Kruger National Park. (3) 12-35, 15-20, 17 ind/1000 km2 [From Table 1] | 1.67, 0.77, 1.2-3.5 | | (1) Maddock and Mills 1994, (2) Marnewick et al. 2014, (3) Fuller et al. 1992 | 2 |
| *Lycaon pictus* | Lake Mutirikwi Recreational Park, Zimbabwe | *There are no large predators in the Park apart from the indigenous side-striped jackal (Canis adustus). Pythons (Python sebae), and crocodiles (Crocodylus niloticus) which are numerous in the lake, may remove a few animals although very few such records exist.* | 0.00 | | Condy 1973 | 0 |

| Predator species scientific name (reference) | Study site name (reference) | Quote | Density (number of individuals /100 km$^2$) | Study site (if different from reference) | Reference | Predator density qualitative index |
|---|---|---|---|---|---|---|
| *Lycaon pictus* | Lewa Wildlife Conservancy, Kenya | Presence not reported on camera trap images | 0.00 | | Sargent 2016 | 0 |
| *Lycaon pictus* | Masai Mara National Reserve, Kenya | 22-35 indiv/1000 km2 [From Table 1] | 2.20-3.50 | | Fuller et al. 1992 | 3 |
| *Lycaon pictus* | Mountain Zebra National Park, South Africa | [species not reported] | 0.00 | | https://www.sanparks.org/ | 0 |
| *Lycaon pictus* | Ngorongoro Crater, Tanzania | *Records for wild dogs Lycaon pictus are rare and they have been described as being present only intermittently (Homewood & Rodgers 1991). Fosbrooke (1972) describes the movements of wild dogs into and out of the crater between the years 1961 and 1966, and the species becoming completely absent in 1962–63. Estes and Goddard (1967) and Schaller (1972) also recorded that wild dogs were present in the 1960s. Packer et al. (1991b) state that they had been 'absent for nearly 20 years' and none was recorded by Rees et al. (2006) during censuses in 2003–06.* | NA | | Oates and Rees 2013 | 1 |
| *Lycaon pictus* | North, Namibia | (1) All the above factors have led to the wild dog becoming highly restricted and rare today, where 30 years ago it was fairly common and widespread. Wild dogs continue to suffer under a multitude of pressures and could be regarded as the most endangered large mammal in Namibia today. If the situation with regard to their requirements for long-term survival are not met soon, wild dogs could be extinct in this country within the next 10-20 years. (2) we identified seven individuals (individuals A, C-G, I). | NA | | (1) Hines 1990, (2) Hofmann et al. 2025 | 2 |
| *Lycaon pictus* | Pilanesberg National Park, South Africa | *Nine wild dogs were introduced to Pilanesberg as an artificially bonded pack [but currently, no wild dogs in the study site]* | 0.00-1.57 | | van Dyk and Slotow 2003 | 1 |
| *Lycaon pictus* | Save Valley Conservancy and Imire Game Ranch, Zimbabwe | *SVC is home to the big five [...]; while carnivores are dominated by the lion, spotted hyena, and African wild dog (Lycaon pictus) [26].* | NA | | Makumbe et al. 2022 | 2 |
| *Lycaon pictus* | Serengeti, Tanzania | <10, 10-12, 5-29, 2 indiv/1000 km2 [From Table 1] | 0.20-2.90 | | Fuller et al. 1992 | 2 |
| *Lycaon pictus* | Sinamatella Intensive Protection Zone, Zimbabwe | [same reports as Hwange National Park, Zimbabwe considered for this study site] | 0.34-1.80 | | / | 1 |
| *Lycaon pictus* | Tsavo East National Park, Kenya | *wild dog (111 ± 92)* | 0.81 | Tsavo National Parks | Henschel et al. 2020 | 1 |
| *Lycaon pictus* | Umfolozi Game Reserve, South Africa | [same reports as Hluhluwe Corridor Umfolozi Game Reserve Complex, South Africa considered for this study site] | 3.75-9.90 | | / | 3 |
| *Lycaon pictus* | Zambia | *African wild dogs, primarily in one pack, were present until 2014 when they were locally extirpated, likely due to rabies (M'soka et al. 2016), and remained absent during this study.* | 0.00 | Western Zambia's Greater Liuwa Ecosystem | Martens et al. 2025 | 0 |
| *Lycaon pictus* | Ziwa Rhino Sanctuary, Uganda | [species not reported] | 0.00 | | https://ziwarhinoandwildliferanch.com/about- | 0 |

| Predator species scientific name (reference) | Study site name (reference) | Quote | Density (number of individuals /100 km$^2$) | Study site (if different from reference) | Reference | Predator density qualitative index |
|---|---|---|---|---|---|---|
| | | | | | ziwa/wildlife-on-the-ranch/ | |
| *Panthera leo* | Addo Elephant National Park, South Africa | *(1) This is probably because of the low density of each (0.04 lions and 0.07 hyaenas km-2), reducing the likelihood of competitive or agonistic interactions. [estimate seems overestimated] (2) Six lions (four males and two females) were reintroduced to the Main Camp section of the park in 2003 and 2004 (Hayward et al. 2007a, c)* | 0.37 | | (1) Hayward and Hayward 2006, (2) Hayward et al. 2009 | 1 |
| *Panthera leo* | Etosha National Park and Daan Viljoen Game Reserve and Namib Desert Park and Naukluft Mountain Zebra Park, Namibia | *(1) Between 1983 and 1997, the total lion population in central Etosha fluctuated between 50 and 88 individuals (Fig. 5). The total number of adult and subadult lions, however, decreased over the years and reached an equilibrium of about 42 individuals, which was caused by the decrease of adult females (Figs. 5 and 6). There were three distinctive peaks in population size resulting from a larger number of cubs produced in 1984, 1989 and 1996, which can be related to rainfall: There existed a strong correlation between cubs per female and annual rainfall (Fig. 7). (2) Initially, perimeter fencing and the construction of artificial waterpoints appeared to maximise Etosha's lion population, which increased from approximately 200 to 500 individuals during the 1970s. (3) lion density estimated at 1.6-2 individuals per 100km2* | 0.22-0.40, 0.90-2.25, 1.60-2.00 | | (1) Trinkel 2013, (2) Heydinger and Funston 2022, (3) Stander 1991 | 1 |
| *Panthera leo* | Hluhluwe Corridor Umfolozi Game Reserve Complex, South Africa | *(1) The native HiP population consisted of about 84 lions in 2000 but crashed to only 20 native individuals and their offspring by 2004, corresponding to 32% of the total population (Fig. 1). F1 offspring of translocated and native HiP lions totalled 29 individuals by the end of 2004 (47%), and the translocated lions and their offspring totalled 13 individuals (21%) (Fig. 1). (2) However, the extirpation of the other members of the large carnivore guild in the first half of the 20th century has meant that African lions (Panthera leo), African wild dogs (Lycaon pictus), and cheetahs (Acinonyx jubatus) have all had to be reintroduced (Cromsigt et al. 2017; Trinkel et al. 2008). The estimated numbers of these latter three species in 2018 were 70 [lion] [...] (Roberts 2022).* | 2.08-8.75 | | (1) Trinkel et al. 2008, (2) Roberts et al. 2023 | 2 |
| *Panthera leo* | Hwange National Park, Zimbabwe | *(1) Home range and demographic data suggest that the density of the population in HNP was around 2.7 lions/100 km2. (2) In the study area, the [...] lion density was 3.5 lions/100 km2 (Andrew J. Loveridge, pers. com).* | 2.7-3.5 | | (1) Loveridge et al. 2007, (2) Périquet et al. 2016 | 1 |
| *Panthera leo* | Ithala Game Reserve, South Africa | *IGR has been virtually predator free since its creation in 1972.* | 0.00 | | O'Kane and Macdonald 2016 | 0 |
| *Panthera leo* | Kruger National Park, South Africa | *(1) Disregarding zonal differences, we estimated a population size of 1684 (95% CI: 1617–1751) lions for Kruger as a whole. (2) 9 individual per 100 km2* | 8.64-9.00 | | (1) Ferreira and Funston 2010, (2) Stander 1991 | 2 |
| *Panthera leo* | Lake Mutirikwi Recreational Park, Zimbabwe | *There are no large predators in the Park apart from the indigenous side-striped jackal (Canis adustus). Pythons (Python sebae), and crocodiles (Crocodylus niloticus) which are numerous in the lake, may remove a few animals although very few such records exist.* | 0.00 | | Condy 1973 | 0 |
| *Panthera leo* | Lewa Wildlife Conservancy, Kenya | Presence reported on camera trap images | NA | | Sargent 2016 | 2 |

| Predator species scientific name (reference) | Study site name (reference) | Quote | Density (number of individuals /100 km$^2$) | Study site (if different from reference) | Reference | Predator density qualitative index |
|---|---|---|---|---|---|---|
| *Panthera leo* | Masai Mara National Reserve, Kenya | *(1) Overall posterior mean lion density was estimated to be 17.08 (posterior SD 1.310) lions >1 year old/100 km2, and the sex ratio was estimated at 2.2 females to 1 male. (2) lion density was anomalously 8.0 times lower in the ranches (0.046 lions/km2) than in the reserve (0.369 lions/km2). Lion and hyena densities and prey biomass did not differ between June 1991 (5172.273 kg/km2) and June 2003 (5472 kg/km2) in the reserve [...]. Lions never responded to playbacks in the ranches, so the potential shift in lion behavioural response for different land use zones is another potential explanation for the patterns found here. (3) 0.3 individual per 100 km2* | 17.08, 4.60-36.90, 0.30 | | (1) Eliot and Gopalaswamy 2016, (2) Ogutu at al. 2005, (3) Stander 1991 | 3 |
| *Panthera leo* | Mountain Zebra National Park, South Africa | *Lions Panthera leo have been reintroduced into one privately-owned and three formally protected areas which have CMZ, namely Kuzuko, Karoo NP and MZNP i.e. the two largest CMZ populations are exposed to lions. [...] At least 13 CMZ were killed by the two lion in MZNP over a two year period (2013-2014) and in Karoo NP, lion showed a preference for CMZ during the initial post-release period when their movements were concentrated within the high CMZ density area on the mountain tops (their prey preference has since shifted to kudu, red hartebeest and gemsbok; personal communication with SANParks).* | 0.70 | | Hrabar and Kerley 2015 | 1 |
| *Panthera leo* | Ngorongoro Crater, Tanzania | *(1) The population of adult lions in the Crater increased from approximately 20 in the late 1960s to about 40 in the late 1970s, remained at this level until the early 1990s, and then declined to about 25 by 1996 (Packer et al. 1991, pers. obs.). (2) In 1962, the population declined from over 60 lions to just 15 individuals (Fosbrooke 1963). It subsequently recovered, reaching a peak of 124 in 1983, but then entered a gradual decline phase. Closely spaced die-offs in 1994, 1997 and 2001 have meant that the population has not had sufficient time to recover in between, resulting in fewer than 60 lions in the crater since 1994 (Kissui & Packer 2004). (3) 27 individual per 100 km2* | 7.69-15.38, 5.77-47.69, 27.00 | | (1) Höner et al. 2005, (2) Oates and Rees 2013, (3) Stander 1991 | 3 |
| *Panthera leo* | North, Namibia | *The population is estimated between 57 and 60 individual adult lions and 14 cubs; this represents an inferred decrease of 46–60% over the past five years. At 0.11–0.12 lions/100 km2, this is the lowest recorded density for a free-ranging, self-sustaining lion population in Africa. Thirty-six female and 21 male lions were found during the survey, yielding a sex ratio of 1 female: 0.58 male.* | 0.11-0.12 | | Heydinger et al. 2024 | 1 |
| *Panthera leo* | Pilanesberg National Park, South Africa | *Lions were introduced in 1993 and presently number 60 individuals.The Pilanesberg lion population consisted of 13 adult males, 13 adult females and 15 cubs (<1.5 years old), and 10 subadults (<3 years old).* | 10.49 | | van Dyk and Slotow 2003 | 2 |
| *Panthera leo* | Save Valley Conservancy and Imire Game Ranch, Zimbabwe | *Results showed that lions (Panthera leo), spotted hyenas (Crocuta crocuta), elephants (Loxodonta africana), and Nile crocodiles (Crocodylus niloticus) were the major animals involved in the conflict.* | NA | | Makumbe et al. 2022 | 2 |
| *Panthera leo* | Serengeti, Tanzania | (1) between 0.01 and 0.45 individuals per km2 [From Figure 3]. (2) 7.9-9.4 individual per 100 km2 | 1.00-45.00, 7.90-9.40 | | (1) Durant et al. 2011, (2) Stander 1991 | 3 |
| *Panthera leo* | Sinamatella Intensive Protection Zone, Zimbabwe | [same reports as Hwange National Park, Zimbabwe considered for this study site] | 2.7-3.5 | | / | 1 |
| *Panthera leo* | Tsavo East National Park, Kenya | *lion (706 ± 201)* | 5.14 | Tsavo National Parks | Henschel et al. 2020 | 2 |
| *Panthera leo* | Umfolozi Game Reserve, South Africa | [same reports as Hluhluwe Corridor Umfolozi Game Reserve Complex, South Africa considered for this study site] | 2.08-8.75 | | / | 2 |

| **Predator species scientific name (reference)** | **Study site name (reference)** | **Quote** | **Density (number of individuals /100 km$^2$)** | **Study site (if different from reference)** | **Reference** | **Predator density qualitative index** |
|---|---|---|---|---|---|---|
| *Panthera leo* | Zambia | *(1) On the other hand, Chardonnet [3] utilized a finer-scale resolution to produce an estimate of 3575 lions in Zambia, but for some areas their calculations relied on lion densities recorded nearly 40 years ago [13]. Given the global [14] [15] and continental [16] [17] declines of large carnivore populations over the same period, it is unlikely that historic numbers accurately describe current status. (2) lions were nearly absent from the system, save for one lioness, for several years prior to this study. After subsequent reintroductions of two, two, and one lion in 2009, 2011, and 2016 respectively, the population increased to 10 individuals by 2019. The near absence of lions (< 0.3 individuals/100 km2, more than an order of magnitude less than other wildebeest-dominated ecosystems; Creel, unpublished data; Durant et al. 2011; Elliot and Gopalaswamy 2017) in a system with a large prey base has largely released the hyena population from lion competition and predation, allowing the hyena to become the apex predator. (3) the African lion, was severely reduced to 3–5 animals.* | <0.30, 0.12 | (2) Western Zambia's Greater Liuwa Ecosystem, (3) Liuwa Plain National Park | (1) Chomba et al. 2014, (2) Martens et al. 2025, (3) M'soka et al. 2016 | 1 |
| *Panthera leo* | Ziwa Rhino Sanctuary, Uganda | *it has no predators such as lions* | 0.00 | | Patton and Genade 2021 | 0 |
| *Panthera pardus* | Addo Elephant National Park, South Africa | *a male leopard [...] reintroduced to the Main Camp section of the park in 2003 and 2004 (Hayward et al. 2007a, c)* | 0.06 | | Hayward et al. 2009 | 1 |
| *Panthera pardus* | Etosha National Park and Daan Viljoen Game Reserve and Namib Desert Park and Naukluft Mountain Zebra Park, Namibia | *We identified [...] 15 individual [...] leopards* | 0.07 | | Keja et al. 2025 | 1 |
| *Panthera pardus* | Hluhluwe Corridor Umfolozi Game Reserve Complex, South Africa | *In 2018, leopards were believed to be present at a density of 3.3 ± 0.8 leopards per 100 km2 (Mann et al. 2019a,b).* | 3.30 | | Roberts et al. 2023 | 1 |
| *Panthera pardus* | Hwange National Park, Zimbabwe | Densities between 0.01 and 0.02 individuals per km2 [from Table 1] | 1.00-2.00 | | Loveridge et al. 2022 | 1 |
| *Panthera pardus* | Ithala Game Reserve, South Africa | *IGR has been virtually predator free since its creation in 1972.* | 0.00 | | O'Kane and Macdonald 2016 | 0 |
| *Panthera pardus* | Kruger National Park, South Africa | *Density varied substantially among sites, ranging from 2.6 ± 0.6 leopards/100 km2 to 13.2 ± 2.6 leopards/100 km2* | 2.60-13.20 | | Smyth et al. 2024 | 2 |
| *Panthera pardus* | Lake Mutirikwi Recreational Park, Zimbabwe | *There are no large predators in the Park apart from the indigenous side-striped jackal (Canis adustus). Pythons (Python sebae), and crocodiles (Crocodylus niloticus) which are numerous in the lake, may remove a few animals although very few such records exist.* | 0.00 | | Condy 1973 | 0 |
| *Panthera pardus* | Lewa Wildlife Conservancy, Kenya | Presence not reported on camera trap images | 0.00 | | Sargent 2016 | 0 |
| *Panthera pardus* | Masai Mara National Reserve, Kenya | *We recorded 725 leopard images and estimated population density at 1.90 ± 0.56 individuals 100 km2−1, relatively low compared to other areas and only slightly higher than previous MME estimates of cheetah, an ecologically subordinate competitor* | 1.90 | | Hills et al. 2025 | 1 |

| Predator species scientific name (reference) | Study site name (reference) | Quote | Density (number of individuals /100 km$^2$) | Study site (if different from reference) | Reference | Predator density qualitative index |
|---|---|---|---|---|---|---|
| *Panthera pardus* | Mountain Zebra National Park, South Africa | [species not reported] | 0.00 | | https://www.sanparks.org/ | 0 |
| *Panthera pardus* | Ngorongoro Crater, Tanzania | *Too few records of leopards exist to draw any conclusions about their history in the crater.* | NA | | Oates and Rees 2013 | 2 |
| *Panthera pardus* | North, Namibia | *(1) With an effort of 2430 and 2074 camera trap nights in the KNP and LHR, respectively, 11 adult female and six adult male leopards were identified in the KNP, whilst only one adult female leopard was detected once in the LHR. For the KNP, a maximum likelihood approach (using the package SECR) revealed a density estimate of 2.74 leopards/100 km2, whereas a Bayesian approach (using the package SPACECAP) revealed a density estimate of 1.83 leopards/100 km2. For the LHR, no density estimate could be determined and it is suggested that the leopard density in such an arid environment is low. (2) Using the best-fitting model (heterogeneity – (M(h))within program CAPTURE, we calculated that the leopard density (no. / 100 km2) was significantly higher (Z-score = 2.4; P = 0.02) in farmland areas (3.6, 95%CI = 3.0–8.0) than within the Park (1.0, 0.8–1.5; Table 2). The ratio of photographed to nonphotographed radio-marked leopards in the farmlands (2 / 3; 0.67) was similar to the ratio of total leopards photographed in the farmlands to the calculated estimated total there (10 / 13; 0.77).* | 1.83, 1.00-3.60 | Khaudum National Park and Lower Hoanib River | (1) Portas et al. 2022, (2) Stein et al. 2011 | 1 |
| *Panthera pardus* | Pilanesberg National Park, South Africa | *Other large predators include leopards (Panthera pardus) (estimated population of 40–60 individuals)* | 8.74 | | van Dyk and Slotow 2003 | 3 |
| *Panthera pardus* | Save Valley Conservancy and Imire Game Ranch, Zimbabwe | *SVC is home to the big five; buffalo (Syncerus caffer), elephant, lion, leopard, and black rhino* | NA | | Makumbe et al. 2022 | 2 |
| *Panthera pardus* | Serengeti, Tanzania | *(1) Other studies include densities of 6.81 ± 1.24 100 km2−1 in the Serengeti National Park, Tanzania, which is contiguous with the MME (Allen et al. 2020). (2) We estimated leopard densities, at 5.41 (95% CrI = 2.23–9.26) and 5.72 (95% CrI = 2.44–9.55) individuals/100 km2, in the dry and wet season, respectively, which confirmed Serengeti National Park as one of the strongholds of this species in Africa.* | 6.81, 5.41-5.72 | | (1) Hills et al. 2025, (2) Allen et al. 2020 | 2 |
| *Panthera pardus* | Sinamatella Intensive Protection Zone, Zimbabwe | [same reports as Hwange National Park, Zimbabwe considered for this study site] | 1.00-2.00 | | / | 1 |
| *Panthera pardus* | Tsavo East National Park, Kenya | *leopard (452 ± 98)* | 2.05 | Tsavo National Parks | Henschel et al. 2020 | 1 |
| *Panthera pardus* | Umfolozi Game Reserve, South Africa | [same reports as Hluhluwe Corridor Umfolozi Game Reserve Complex, South Africa considered for this study site] | 3.30 | | / | 1 |
| *Panthera pardus* | Zambia | *(1) Leopards are absent. (2) Population estimates resulted in 12 individuals for LNP 2008 and 10 for GMA-A. The selected part inside the GMA-A which is smaller in area, reflected a population density estimate of 4.79 ± 1.16 per 100 km², higher than that recorded in the National Park at 3.36 ± 0.64.* | 0.00, 3.36-4.79 | (1) Western Zambia's Greater Liuwa Ecosystem, (2) Luambe National Park and bordering Game | (1) Martens et al. 2025, (2) Ray 2011 | 1 |

| **Predator species scientific name (reference)** | **Study site name (reference)** | **Quote** | **Density (number of individuals /100 km$^2$)** | **Study site (if different from reference)** | **Reference** | **Predator density qualitative index** |
|---|---|---|---|---|---|---|
| | | | | Management Area | | |
| *Panthera pardus* | Ziwa Rhino Sanctuary, Uganda | [species reported] | NA | | https://ziwarhinoandwildliferanch.com/about-ziwa/wildlife-on-the-ranch/ | 2 |
| *Panthera tigris* | Chang Tang Reserve, China | [species not reported] | 0.00 | | / | 0 |
| *Panthera tigris* | Chitwan National Park, Nepal | *In 2013 tiger density was estimated to be 3.84 per 100 km2 [...] (Dhakal et al., 2014).* | 3.84 | | Dhungana et al. 2017 (from other sources) | 1 |
| *Panthera tigris* | Gobi A National Park, Mongolia | [species not reported] | 0.00 | Tost Local Protected Area | Tumursukh et al. 2016 | 0 |
| *Panthera tigris* | Gunung Leuser National Park, Indonesia | *During the 90-day sessions, sex-specific tiger densities ranged from 0.97 (± 0.37)–1.83 (± 0.59) female and 0.56 (± 0.22)–0.61 (± 0.29) male tigers. During the 180-day sessions, estimates were only negligibly more precise, ranging from 0.94 (± 0.33)–1.76 (± 0.54) female and 0.41 (± 0.15)–0.50 (± 0.22) male tigers. Across both years, densities were 1.42–2.35 tigers/100 km2* | 1.42-2.35 | | Figel et al. 2025 | 1 |
| *Panthera tigris* | Hustai National Park, Mongolia | [Tiger appears to be extinct in Mongolia] | 0.00 | | / | 0 |
| *Panthera tigris* | Takhin Tal, Mongolia | [Tiger appears to be extinct in Mongolia] | 0.00 | | / | 0 |
| *Panthera tigris* | Xinjiang, China | *Caspian tigers (Panthera tigris virgata), whose range once spanned a vast area across Xinjiang in China and central Asia, went extinct in the 1980s due to intense human interference, including deforestation, development, and hunting (Chestin et al., 2017).* | 0.00 | | Zhang et al. 2023 | 0 |
| *Panthera uncia* | Chang Tang Reserve, China | *In addition to the 270 new snow leopard location points collected by the authors, an additional 39 snow leopard locations reported by other observers in the study area were compiled from the three sources discussed in Section 2.2.5, above (Fig. 1, Table 2). These included 22 pre-2007 locations from the unpublished 2007 SLT-WCS-Panthera-SLN range-wide snow leopard distribution map and an additional 10 snow leopard location points contributed during the 2008 Beijing conference, six of which were reported by Joseph Fox, one by George Schaller, and three of which were unattributed. The last source was the snow leopard scat collection location map published by Zhou et al. (2014). This map provided four additional snow leopard scat locations for Shainza County and three for Pelgon County for scat samples collected at an unspecified time between 2006 and 2009.* | NA | | Farrington and Tsering 2020 | 2 |
| *Panthera uncia* | Chitwan National Park, Nepal | *Leopard population in Chitwan National Park, its buffer zone and adjoining forests was estimated 107 (95 % CI: 81–144) with density 3.95 (95 % CI: 2.76–5.2) leopards per 100 km2 in 2022. We documented stable leopard population with a slight decline from 58 (37–77) in 2013 to 48 (34–66) in 2022 in the park* | 3.95 | | Shrestha et al. 2025 | 3 |
| *Panthera uncia* | Gobi A National Park, Mongolia | *The adult snow leopard population in our study area during 2012-2013, estimated independently using camera-trap-based mark–recapture methods, was 12-14.* | 0.93 | Tost Local Protected Area | Tumursukh et al. 2016 | 1 |

| Predator species scientific name (reference) | Study site name (reference) | Quote | Density (number of individuals /100 km$^2$) | Study site (if different from reference) | Reference | Predator density qualitative index |
|---|---|---|---|---|---|---|
| *Panthera uncia* | Hustai National Park, Mongolia | *Mongolia contains significant populations of many of these globally threatened species, such as the Asiatic wild ass (E. hemionus), Przewalski's horse, Bactrian camel, saiga antelope and snow leopard.* | NA | | Clark et al. 2005 | 2 |
| *Panthera uncia* | Takhin Tal, Mongolia | *SECR analysis resulted in an overall density of 1.31 individuals/100 km2 (1.15%–1.50 95% CI), which was positively correlated with terrain slope* | 1.31 | Altai Mountains | Oberosler et al. 2022 | 2 |
| *Panthera uncia* | Xinjiang, China | *In Xinjiang, Schaller (1988a) estimated that there were no more than 750 individuals within an area of 170 000 km² suitable habitat. In Qinghai, he indicated that their numbers could be in the order of 650 across the province's 65 000 km² range, based on an estimated density of one snow leopard per 100 km2 (Schaller et al. 1988b). More recent estimates based on systematic sign and camera trap surveys (without the use of mark–recapture analysis) in Xinjiang have produced estimates of 2–5 per 100 km2 (Ma et al. 2006; Xu et al. 2011a,b). Wu (2009) estimated, on the basis of signs and social surveys, that there was a minimum of 7 individuals within a 1900 km2 area in Xinjiang. Peng (2009) estimated that there were 51–78 snow leopard individuals across 9 nature reserves within Ganzi Prefecture, Sichuan. Finally genetic identification from scats was used by Janečka et al. (2008) to identify 1 male snow leopard in Qinghai during a brief 2-day survey. Similarly, Zhou et al. (2014) applied genetic methods to identify 48 snow leopard individuals across Xizang, Qinghai and Gansu. Using capture–recapture methods, McCarthy et al. (2008) estimated a density of 0.74 individuals per 100 km2 in the Tien Shan mountain range.* | 0.44, 2.00-5.00, 0.37 | | Alexander et al. 2016 | 3 |

Table S3.3: Density estimates for snow leopard (*Panthera uncia*) and Nile crocodile (*Crocodylus niloticus*) extracted from the literature.

| **Predator scientific name (reference)** | **Locality** | **Country** | **Density** | **Reference** |
|---|---|---|---|---|
| *Crocodylus niloticus* | Akagera National Park | Rwanda | 31.11 | Combrink et al. 2019 |
| *Crocodylus niloticus* | Amatigulu Nature Reserve | South Africa | 75.00 | Combrink et al. 2019 |
| *Crocodylus niloticus* | Gonarezhou National Park | Zimbabwe | 06.08 | Combrink et al. 2019 |
| *Crocodylus niloticus* | Hluhluwe Corridor Umfolozi Game Reserve Complex | South Africa | 60.00 | Combrink et al. 2019 |
| *Crocodylus niloticus* | Kruger National Park | South Africa | 17.07 | Combrink et al. 2019 |
| *Crocodylus niloticus* | Limpopo Province | South Africa | 0.71 | Combrink et al. 2019 |
| *Crocodylus niloticus* | Liwonde National Park | Malawi | 123.36 | Combrink et al. 2019 |
| *Crocodylus niloticus* | Mamili National Park | Namibia | 415.00 | Combrink et al. 2019 |
| *Crocodylus niloticus* | Maputo Special Reserve | Mozambique | 1.60 | Combrink et al. 2019 |
| *Crocodylus niloticus* | Mpumalanga Province | South Africa | 0.48 | Combrink et al. 2019 |
| *Crocodylus niloticus* | Murchison Falls National Park | Uganda | 11.83 | Combrink et al. 2019 |
| *Crocodylus niloticus* | Okavango Delta | Botswana | 6.18 | Combrink et al. 2019 |
| *Crocodylus niloticus* | Pilanesberg National Park | South Africa | 4.53 | Power and Verburgt 2014 |
| *Crocodylus niloticus* | Save Valley Conservancy and Imire Game Ranch | Zimbabwe | 9.19 | Combrink et al. 2019 |
| *Crocodylus niloticus* | Sengwa Wildlife Research Area | Zimbabwe | 10.99 | Combrink et al. 2019 |
| *Crocodylus niloticus* | Lower Zambesi Region | Zimbabwe and Zambia | 44.03 | Wallace et al. 2013 |
| *Crocodylus niloticus* | Ndumo Game Reserve | South Africa | 846.00 | Calverley and Downs 2014 |
| *Panthera uncia* | Altai Mountains | Mongolia | 1.31 | Oberosler et al. 2022 |
| *Panthera uncia* | Chitwan National Park | Nepal | 3.95 | Shrestha et al. 2025 |
| *Panthera uncia* | Tost Local Protected Area | Mongolia | 0.93 | Tumursukh et al. 2016 |
| *Panthera uncia* | Qinghai | China | 1.00 | Alexander et al. 2016 |
| *Panthera uncia* | Tien Shan Mountain Range | China | 0.74 | Alexander et al. 2016 |
| *Panthera uncia* | Xinjiang | China | 0.44 | Alexander et al. 2016 |
| *Panthera uncia* | Xinjiang | China | 0.37 | Alexander et al. 2016 |
| *Panthera uncia* | Xinjiang | China | 3.50 | Alexander et al. 2016 |

Density estimates are reported as the number of individuals per 100 $km^2$.

Supporting information 4: Correlations between predictors investigated for their effect on birth synchrony in Perissodactyla.

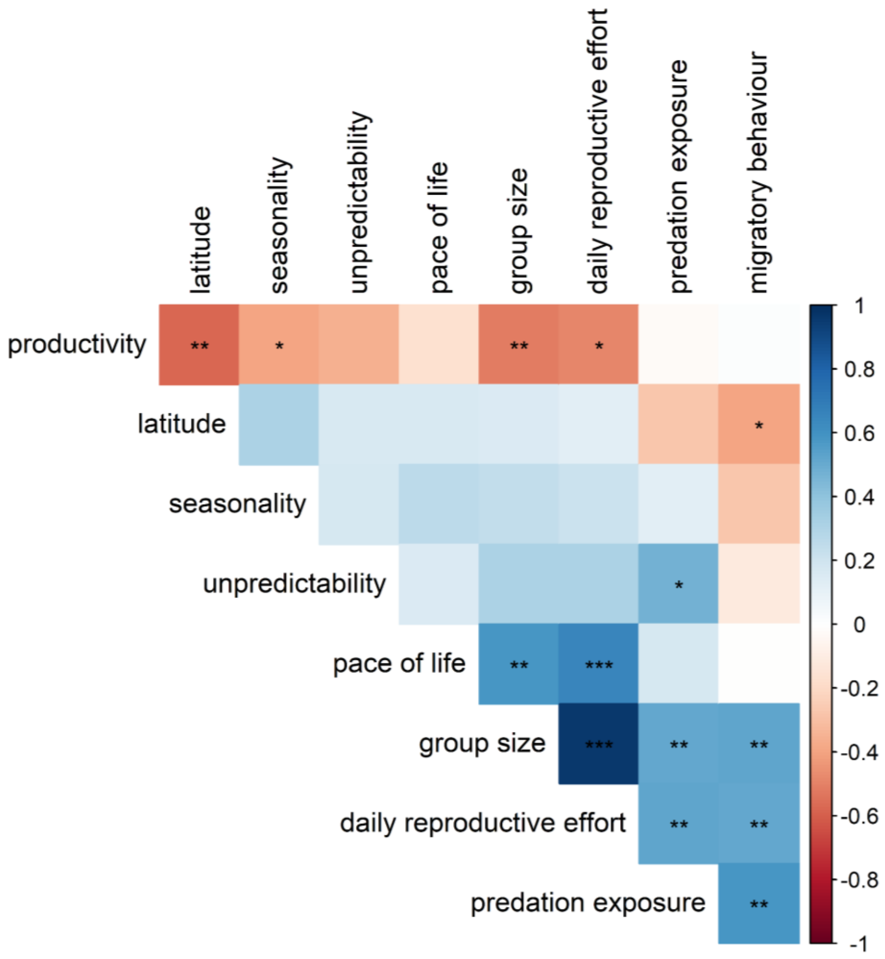


Figure 4.1: Correlations (Spearman's rank correlation $\rho$) between the nine predictors (latitude, environmental seasonality, environmental productivity, environmental unpredictability, intensity of daily reproductive effort, migratory behaviour, pace of life, group size and predation exposure) investigated for their effect on birth synchrony in 27 populations of Perissodactyla species ($n$ = 10). Colours show the strength of the correlation ($\rho$, from the highest negative correlation in red to the highest positive correlation in blue), stars indicate the level of significance of the correlation to the threshold α (***: α = 0.001, **: α = 0.01,*: α = 0.05).

References supporting information

Abdibek, A.E., Abayeva, K.T., Dosmanbetov, D.A., Baibatshanov, M.K., Khamchukova, A.M., Narbayev, S. et al. (2025). Development of technology for relocating kulans to preserve their population and restore the biodiversity of their habitat. Brazilian Journal of Biology, 85, e291458.

Alexander, J.S., Zhang, C., Shi, K. and Riordan, P. (2016). A spotlight on snow leopard conservation in China. Integrative Zoology, 11(4), 308-321.

Alibhai, S.K., Jewell, Z.C. and Towindo, S.S. (2001). Effects of immobilization on fertility in female black rhino (*Diceros bicornis*). Journal of Zoology, 253(3), 333-345.

Allen, M.L., Wang, S., Olson, L.O., Li, Q. and Krofel, M. (2020). Counting cats for conservation: seasonal estimates of leopard density and drivers of distribution in the Serengeti. Biodiversity and Conservation, 29(13), 3591-3608.

Annear, E., Minnie, L., Andrew, K. and Kerley, G.I. (2023). Can smaller predators expand their prey base through killing juveniles? The influence of prey demography and season on prey selection for cheetahs and lions. Oecologia, 201(3), 649-660.

Ansell, W.F.H. (1960). The breeding of some larger mammals in Northern Rhodesia. Proceedings of the Zoological Society of London, 134(2), 251-274.

Balajeid Lyngdoh, S., Habib, B. and Shrotriya, S. (2020). Dietary spectrum in Himalayan wolves: Comparative analysis of prey choice in conspecifics across high-elevation rangelands of Asia. Journal of Zoology, 310(1), 24-33.

Boyd, L.E. (1991). The behaviour of Przewalski's horses and its importance to their management. Applied Animal Behaviour Science, 29, 301-318.

Boyd, L. and Houpt, K.A. (1994). Activity Patterns. In Boyd, L. and Houpt, K. A. (Eds.), Przewalski's horse: the history and biology of an endangered species (pp. 195-227). State University of New York Press.

Broekhuis, F. and Gopalaswamy, A.M. (2016). Counting cats: Spatially explicit population estimates of cheetah (*Acinonyx jubatus*) using unstructured sampling data. PLoS one, 11(5), e0153875.

Calverley, P.M. and Downs, C.T. (2014). Population status of Nile crocodiles in Ndumo Game Reserve, KwaZulu-Natal, South Africa (1971–2012). Herpetologica, 70(4), 417-425.

Chen, J., Weng, Q., Chao, J., Hu, D. and Taya, K. (2008). Reproduction and development of the released Przewalski's horses (*Equus przewalskii*) in Xinjiang, China. Journal of equine science, 19(1), 1-7.

Chomba, C., Senzota, R., Chabwela, H. and Nyirenda, V. (2014). Lion hunting and trophy quality records in Zambia for the period 1967-2000: will the trends in trophy size drop as lion population declines? Open Journal of Ecology, 4(4), 182.

Clark, E.L., Ocock, J.F., King, S.R. and Baillie, J.E. (2005). Proceedings of the Mongolian Biodiversity Databank Workshop: Assessing the Conservation Status of Mongolian Mammals and Fishes: II–Mammals: Assessment Results and Threats. Mongolian Journal of Biological Sciences, 3(2), 17-27.

Clements, H.S., Knight, M., Jones, P. and Balfour, D. (2020). Private rhino conservation: Diverse strategies adopted in response to the poaching crisis. Conservation Letters 13, e12741.

Combrink, X., Lippai, C. and Fergusson, R. (2019). Nile Crocodile Crocodylus niloticus. In Manolis, S. C. and Stevenson, C. (Eds.), Crocodiles: Status Survey and Conservation Action Plan (pp. 1-28). IUCN.

Condy, P.R. (1973). The population status, social behaviour, and daily activity pattern of the white rhinoceros (*Ceratotherium simum simum*) in Kyle National Park, Rhodesia. [MSc dissertation, University of Rhodesia].

Creel, S. and Creel, N.M. (1996). Limitation of African wild dogs by competition with larger carnivores. Conservation Biology, 10(2), 526-538.

Davidson, Z., Dupuis-Desormeaux, M., Dheer, A., Pratt, L., Preston, E., Gilicho, S. et al. (2019). Borrowing from Peter to pay Paul: managing threatened predators of endangered and declining prey species. PeerJ, 7, e7916.

Davis, R.S., Overton, E.K., Prugnolle, F., Rougeron, V., Sievert, O. and Venter, J.A. (2024). Baboons (*Papio* spp.) as a potentially underreported source of food loss and kleptoparasitism of cheetah (*Acinonyx jubatus*) kills. Food Webs, 38, e00331.

Dhungana, R., Savini, T., Karki, J.B., Dhakal, M., Lamichhane, B.R. and Bumrungsri, S. (2018). Living with tigers *Panthera tigris*: patterns, correlates, and contexts of human–tiger conflict in Chitwan National Park, Nepal. Oryx, 52(1), 55-65.

Dinerstein, E. and Price, L. (1991). Demography and habitat use by greater one-horned rhinoceros in Nepal. The Journal of Wildlife Management, 55(3), 401-411.

Dorj, U. and Namkhai, B. (2013). Reproduction and mortality of re-introduced Przewalski's horse *Equus przewalskii* in Hustai National Park, Mongolia. Journal of Life Sciences, 7(6), 623.

Durant, S.M., Craft, M.E., Hilborn, R., Bashir, S., Hando, J. and Thomas, L. (2011). Long-term trends in carnivore abundance using distance sampling in Serengeti National Park, Tanzania. Journal of Applied Ecology, 48(6), 1490-1500.

Elliot, N.B. and Gopalaswamy, A.M. (2017). Toward accurate and precise estimates of lion density. Conservation Biology, 31(4), 934-943.

Fairall, N. (1968). The reproductive seasons of some mammals in the Kruger National Park. African Zoology, 3(2), 189-210.

Farrington, J.D. and Tsering, D. (2020). Snow leopard distribution in the Chang Tang region of Tibet, China. Global Ecology and Conservation, 23, e01044.

Feh, C., Boldsukh, T. and Tourenq, C. (1994). Are family groups in equids a response to cooperative hunting by predators? The case of Mongolian kulans (*Equus hemionus luteus* Matschie). Revue d'écologie, 49(1), 11-20.

Feh, C., Munkhtuya, B., Enkhbold, S. and Sukhbaatar, T. (2001). Ecology and social structure of the Gobi khulan *Equus hemionus* subsp. in the Gobi B National Park, Mongolia. Biological conservation, 101(1), 51-61.

Ferreira, S.M. and Funston, P.J. (2010). Estimating lion population variables: prey and disease effects in Kruger National Park, South Africa. Wildlife Research, 37(3), 194-206.

Ferreira, S.M. and Funston, P.J. (2016). Population estimates of spotted hyaenas in the Kruger National Park, South Africa. African Journal of Wildlife Research, 46(2), 61-70.

Ferreira, S.M. and Pienaar, D. (2011). Degradation of the crocodile population in the Olifants River gorge of Kruger National Park, South Africa. Aquatic Conservation: Marine and Freshwater Ecosystems, 21(2), 155-164.

Figel, J.J., Akbar, M.I., Khairi, K., Darmansyah, Pian, M., Supiyandi, *et al*. (2025). Sumatran tiger density estimates in the Leuser Ecosystem, Sumatra, Indonesia. Frontiers in Conservation Science, 6, 1691233.

Fouché, J., Reilly, B.K., de Crom, E.P., Baeumchen, Y.K. and Forberger, S. (2020). Density estimates of spotted hyaenas (*Crocuta crocuta*) on arid farmlands of Namibia. African Journal of Ecology, 58(5), 3-5.

Freeman, E.W., Meyer, J.M., Bird, J., Adendorff, J., Schulte, B.A. and Santymire, R.M. (2014). Impacts of environmental pressures on the reproductive physiology of subpopulations of black rhinoceros (*Diceros bicornis bicornis*) in Addo Elephant National Park, South Africa. Conservation Physiology, 2(1), cot034.

Fuller, T.K., Kat, P.W., Bulger, J.B., Maddock, A.H., Ginsberg, J.R., Burrows, R. et al. (1992). Population dynamics of African wild dogs. In McCullough, D.R. and Barrett, R.H. (Eds.), Wildlife 2001: Populations (pp. 1125-1139). Springer Dordrecht.

Garnier, J.N., Holt, W.V. and Watson, P.F. (2002). Non-invasive assessment of oestrous cycles and evaluation of reproductive seasonality in the female wild black rhinoceros (*Diceros bicornis minor*). Reproduction, 123(6), 877-889.

Goddard, J. (1967). Home range, behaviour, and recruitment rates of two black rhinoceros populations. African Journal of Ecology, 5(1), 133-150.

Graf, J.A., Somers, M.J., Gunther, M.S. and Slotow, R. (2009). Heterogeneity in the density of spotted hyaenas in Hluhluwe-iMfolozi Park, South Africa. Acta Theriologica, 54(4), 333-343.

Greaver, C., Ferreira, S. and Slotow, R. (2014). Density-dependent regulation of the critically endangered black rhinoceros population in Ithala Game Reserve, South Africa. Austral Ecology, 39(4), 437-447.

Hall-Martin, A.J. and Penzhorn, L. (1977). Behaviour and recruitment of translocated black rhinoceros *Diceros bicornis*. Koedoe, 20(1), 147-162.

Hayward, M.W. (2006). Prey preferences of the spotted hyaena (*Crocuta crocuta*) and degree of dietary overlap with the lion (*Panthera leo*). Journal of Zoology, 270(4), 606-614.

Hayward, M.W. and Hayward, G.J. (2006). Activity patterns of reintroduced lion *Panthera leo* and spotted hyaena *Crocuta crocuta* in the Addo Elephant National Park, South Africa. African journal of ecology, 45(2), 135-141.

Hayward, M.W. and Kerley, G.I. (2005). Prey preferences of the lion (*Panthera leo*). Journal of zoology, 267(3), 309-322.

Hayward, M.W., Hayward, G.J., Druce, D.J. and Kerley, G.I. (2009). Do fences constrain predator movements on an evolutionary scale? Home range, food intake and movement patterns of large predators reintroduced to Addo Elephant National Park, South Africa. Biodiversity and Conservation, 18(4), 887-904.

Hayward, M.W., Henschel, P., O'Brien, J., Hofmeyr, M., Balme, G. and Kerley, G.I. (2006a). Prey preferences of the leopard (*Panthera pardus*). Journal of Zoology, 270(2), 298-313.

Hayward, M.W., Hofmeyr, M., O'brien, J. and Kerley, G.I. (2006b). Prey preferences of the cheetah (*Acinonyx jubatus*) (Felidae: Carnivora): morphological limitations or the need to capture rapidly consumable prey before kleptoparasites arrive? Journal of Zoology, 270(4), 615-627.

Hayward, M.W., Jędrzejewski, W. and Jedrzejewska, B. (2012). Prey preferences of the tiger *Panthera tigris*. Journal of Zoology, 286(3), 221-231.

Hayward, M.W., O'Brien, J., Hofmeyr, M. and Kerley, G.I. (2006c). Prey preferences of the African wild dog *Lycaon pictus* (Canidae: Carnivora): ecological requirements for conservation. Journal of Mammalogy, 87(6), 1122-1131.

Hazarika, B.C. and Saikia, P.K. (2010). A study on the behaviour of great Indian one-horned rhino (*Rhinoceros unicornis* Linn.) in the Rajiv Gandhi Orang National Park, Assam, India. NeBIO, 1(2), 62-74.

Henschel, P., Petracca, L.S., Ferreira, S.M., Ekwanga, S., Ryan, S.D. and Frank, L.G. (2020). Census and distribution of large carnivores in the Tsavo national parks, a critical east African wildlife corridor. African Journal of Ecology, 58(3), 383-398.

Hering-Hagenbeck and Prahl (n.d.). Persian Onager (Equus hemionus onager) EEP. Deutscher-Wildgehege-Verband. Available from: https://www.wildgehege-verband.de/upload/FPDF/Persian_Onager.pdf. [Technical report].

Heydinger, J., Packer, C. and Funston, P. (2022). The historical effects of infrastructure development on the lion population of Etosha National Park, Namibia. Namibian Journal of Environment 6 A: 22-36.

Heydinger, J., Muzuma, U. and Packer, C. (2024). First systematic population survey of the desert-adapted lions, Northwest Namibia. African Journal of Ecology, 62(2), e13266.

Hills, E., Penny, S., Chelysheva, E., Omondi, P., Ngene, S., Giordano, A.J. and Tolhurst, B.A. (2025). Leopard (*Panthera pardus*) Density and the Impact of Spotted Hyaena (*Crocuta crocuta*) Occurrence on Leopard Presence in the Maasai Mara Ecosystem, Kenya. African Journal of Ecology, 63(2), e70025.

Hines, C.J.H. (1990). Past and present distribution and status of the wild dog *Lycaon pictus* in Namibia. Madoqua, 17(1), 31-36.

Hitchins, P.M. and Anderson, J. (1983). Reproduction, population, characteristics and management of the black rhinoceros *Diceros bicornis minor* in the Hluhluwe/Corridor/Umfolozi Game Reserve Complex. South African Journal of Wildlife Research, 13(3), 78-85.

Hoesli, T., Nikowitz, T., Walzer, C. and Kaczensky, P. (2009). Monitoring of agonistic behaviour and foal mortality in free-ranging Przewalski's horse harems in the Mongolian Gobi. Equus, 113-138.

Hofmann, T., Verschueren, S., Shihepo, T., Cristescu, B., Anderson, N., Le Roux, N. et al. (2025). Detection dog survey detects African wild dog presence and a shared marking site. Ecology and Evolution, 15(7), e71703.

Höner, O.P., Wachter, B., East, M.L., Runyoro, V.A. and Hofer, H. (2005). The effect of prey abundance and foraging tactics on the population dynamics of a social, territorial carnivore, the spotted hyena. Oikos, 108(3), 544-554.

Hrabar, H. and Du Toit, J.T. (2005). Dynamics of a protected black rhino (*Diceros bicornis*) population: Pilanesberg National Park, South Africa. Animal Conservation, 8(3), 259-267.

Hrabar, H. and Kerley, G. (2015). Cape Mountain Zebra 2014/15 Status Report, Report 63. Centre for African Conservation Ecology. [Technical report].

Hutchins, M. and Kreger, M.D. (2006). Rhinoceros behaviour: implications for captive management and conservation. International Zoo Yearbook, 40(1), 150-173.

Jacobs, J. (1974). Quantitative measurement of food selection: a modification of the forage ratio and Ivlev's electivity index. Oecologia, 14(4), 413-417.

Joubert, E. (1974). Size and growth as shown by pre-and post-natal development of the Hartmann zebra *Equus zebra hartmannae*. Madoqua, 1(8), 55-58.

Kaczensky, P., Enkhsaikhan, N., Ganbaatar, O. and Walzer, C. (2008). The Great Gobi B Strictly Protected Area in Mongolia-refuge or sink for wolves *Canis lupus* in the Gobi. Wildlife Biology, 14(4), 444-456.

Kamilar, J.M., Bribiescas, R.G. and Bradley, B.J. (2010). Is group size related to longevity in mammals? Biology letters, 6(6), 736-739.

Keja, M., Hauptfleisch, M.L., Nzuma, T.M., Beasley, J., Cloete, C. and Périquet-Pearce, S. (2025). A citizen science survey of cheetahs and leopards in Etosha National Park. African Journal of Wildlife Research, 55, 455-457.

King, S.R.B. (2012). Behavioural ecology of Przewalski horses (*Equus przewalskii*) reintroduced to Hustai National Park, Mongolia. [Doctoral dissertation, Queen Mary University of London].

Klingel, H. (1965). Notes on the biology of the plains zebra *Equus quagga boehmi* Matschie. African Journal of Ecology, 3(1), 86-88.

Klingel, H. (1969). The social organisation and population ecology of the plains zebra (*Equus quagga*). African Zoology, 4(2), 249-263.

Kuntz, R., Kubalek, C., Ruf, T., Tataruch, F. and Arnold, W. (2006). Seasonal adjustment of energy budget in a large wild mammal, the Przewalski horse (*Equus ferus przewalskii*) I. Energy intake. Journal of Experimental Biology, 209(22), 4557-4565.

Laurie, W.A. (1978). The ecology and behaviour of the greater one-horned rhinoceros. [Doctoral dissertation, University of Cambridge].

Leuthold, W. and Leuthold, B.M. (1975). Temporal patterns of reproduction in ungulates of Tsavo East National Park, Kenya. African Journal of Ecology, 13, 159-169.

Loveridge, A.J., Searle, A.W., Murindagomo, F. and Macdonald, D.W. (2007). The impact of sport-hunting on the population dynamics of an African lion population in a protected area. Biological conservation, 134(4), 548-558.

Loveridge, A.J., Sousa, L.L., Seymour-Smith, J.L., Mandisodza-Chikerema, R. and Macdonald, D.W. (2022). Environmental and anthropogenic drivers of African leopard *Panthera pardus* population density. Biological Conservation, 272, 109641.

Lushchekina, A.A., Karimova, T.Y. and Neronov, V.M. (2022). Ungulates of the arid ecosystems from the Red Data Book of the Russian Federation. Arid Ecosystems, 12(4), 432-440.

Lyngdoh, S., Shrotriya, S., Goyal, S.P., Clements, H., Hayward, M.W. and Habib, B. (2014). Prey preferences of the snow leopard (*Panthera uncia*): regional diet specificity holds global significance for conservation. PloS one, 9(2), e88349.

M'soka, J., Creel, S., Becker, M.S. and Droge, E. (2016). Spotted hyaena survival and density in a lion depleted ecosystem: The effects of prey availability, humans and competition between large carnivores in African savannahs. Biological Conservation, 201, 348-355.

Maddock, A.H. and Mills, M.G.L. (1994). Population characteristics of African wild dogs *Lycaon pictus* in the eastern Transvaal lowveld, South Africa, as revealed through photographic records. Biological Conservation, 67(1), 57-62.

Makumbe, P., Mapurazi, S., Jaravani, S. and Matsilele, I. (2022). Human-Wildlife Conflict in Save Valley Conservancy: Residents' Attitude Toward Wildlife Conservation. Scientifica, 2022(1), 2107711.

Marnewick, K., Ferreira, S.M., Grange, S., Watermeyer, J., Maputla, N. and Davies-Mostert, H.T. (2014). Evaluating the status of and African wild dogs *Lycaon pictus* and cheetahs *Acinonyx jubatus* through tourist-based photographic surveys in the Kruger National Park. PloS one, 9(1), e86265.

Martens, S., Creel, S., Becker, M.S., Spong, G., Smit, D., Dröge, E. *et al*. (2025). Long-Term Demography of Spotted Hyena (*Crocuta crocuta*) in a Lion-Depleted but Prey-Rich Ecosystem. Ecology and Evolution, 15(4), e71025.

Miller, D.J. and Schaller, G.B. (1996). Rangelands of the Chang Tang wildlife reserve in Tibet. Rangelands, 18(3), 91-96.

Moehlman, P.D.R. (2002). Equids: Zebras, Asses, and Horses: Status Survey and Conservation Action Plan. IUCN.

Monks, N.J. (1995). The population status, diurnal activity patterns, range and territory size, and habitat use by the white rhinoceros (*Ceratotherium simum*) in Kyle Recreational Park, Zimbabwe [MSc dissertation, University of Kent].

Morin, D.J., Boulanger, J., Bischof, R., Lee, D.C., Ngoprasert, D., Fuller, A.K. et al. (2022). Comparison of methods for estimating density and population trends for low-density Asian bears. Global Ecology and Conservation, 35, e02058.

Mukeka, J.M., Ogutu, J.O., Kanga, E. and Roskaft, E. (2018). Characteristics of human-wildlife conflicts in Kenya: Examples of Tsavo and Maasai Mara regions. Environment and Natural Resources Research, 8(3), 148.

Muntifering, J.R., Guerier, A., Beytell, P. and Stratford, K. (2023). Population parameters, performance and insights into factors influencing the reproduction of the black rhinoceros *Diceros bicornis* in Namibia. Oryx, 57(5), 659-669.

Myhrvold, N.P., Baldridge, E., Chan, B., Sivam, D., Freeman, D.L. and Ernest, S.M. (2015). An amniote life-history database to perform comparative analyses with birds, mammals, and reptiles: Ecological Archives E096-269. Ecology, 96(11), 3109-3109.

Ngana, J.J., Luboobi, L.S. and Kuznetsov, D. (2014). Mathematical model for the population dynamics of the Serengeti ecosystem. Applied and Computational Mathematics, 3(4), 171-176.

Niedziałkowska, M., Hayward, M.W., Borowik, T., Jędrzejewski, W. and Jędrzejewska, B. (2019). A meta-analysis of ungulate predation and prey selection by the brown bear *Ursus arctos* in Eurasia. Mammal Research, 64(1), 1-9.

Nowzari, H., Hemami, M., Karami, M., Kheirkhah Zarkesh, M.M., Riazi, B. and Rubenstein, D. I. (2013). Habitat use by the Persian onager, *Equus hemionus onager* (Perissodactyla: Equidae) in Qatrouyeh National Park, Fars, Iran. Journal of Natural History, 47(43-44), 2795-2814.

O'Kane, C.A. and Macdonald, D.W. (2016). An experimental demonstration that predation influences antelope sex ratios and resource-associated mortality. Basic and applied ecology, 17(4), 370-376.

Oates, L. and Rees, P.A. (2013). The historical ecology of the large mammal populations of Ngorongoro Crater, Tanzania, East Africa. Mammal Review, 43(2), 124-141.

Oberosler, V., Tenan, S., Groff, C., Krofel, M., Augugliaro, C., Munkhtsog, B. and Rovero, F. (2022). First spatially-explicit density estimate for a snow leopard population in the Altai Mountains. Biodiversity and Conservation, 31(1), 261-275.

Ogutu, J.O., Bhola, N. and Reid, R. (2005). The effects of pastoralism and protection on the density and distribution of carnivores and their prey in the Mara ecosystem of Kenya. Journal of Zoology, 265(3), 281-293.

Ogutu, J.O., Piepho, H.P. and Dublin, H.T. (2014). Responses of phenology, synchrony and fecundity of breeding by African ungulates to interannual variation in rainfall. Wildlife Research, 40(8), 698-717.

Owen-Smith, R.N. (1988). Megaherbivores: the influence of very large body size on ecology. Cambridge university press.

Owen-Smith, R.N. (1975). The social ethology of the White Rhinoceros *Ceratotherium simum* (Burchell 1817). Zeitschrift für Tierpsychologie, 38(4), 337-384.

Packer, C., Loveridge, A., Canney, S., Caro, T., Garnett, S., Pfeifer, M. et al. (2013). Conserving large carnivores: dollars and fence. Ecology Letters 16, 635-641.

Paklina, N.V. and van Orden, C. (2007). Territorial Behaviour of Kiang (*Equus kiang* Moorcroft, 1841) in Ladakh (India). Erforschung Biologischer Ressourcen der Mongolei 10, 205-211.

Patton, F. and Genade, A.E. (2021). Behavioural observations of white rhinos at Ziwa Rhino Sanctuary—Analyses from 10 years of data collection. Uganda: Rhino Fund. [Technical report].

Patton, F., Campbell, P.E. and Genade, A. (2022). White rhino ecology: a comparison of two rhino populations (*Ceratotherium simum simum*) in South Africa and Uganda. Pachyderm, 64, 144-148.

Pekor, A., Miller, J.R., Flyman, M.V., Kasiki, S., Kesch, M.K., Miller, S.M. et al. (2019). Fencing Africa's protected areas: Costs, benefits, and management issues. Biological Conservation 229, 67-75.

Penzhorn, B.L. (1985). Reproductive characteristics of a free-ranging population of Cape mountain zebra (*Equus zebra zebra*). Reproduction, 73(1), 51-57.

Périquet, S., Mapendere, C., Revilla, E., Banda, J., Macdonald, D.W., Loveridge, A.J. and Fritz, H. (2016). A potential role for interference competition with lions in den selection and attendance by spotted hyaenas. Mammalian Biology, 81(3), 227-234.

Portas, R., Wachter, B., Beytell, P., Uiseb, K.H., Melzheimer, J. and Edwards, S. (2022). Leopard Panthera pardus camera trap surveys in the arid environments of northern Namibia. Mammalian Biology, 102(4), 1185-1198.

Power, R.J. and Verburgt, L. (2014). The herpetofauna of the North West Province: a literature survey. Department of Economic Development. Environment, Conservation & Tourism, North West Provincial Government, Mahikeng. [Technical report].

Purchase, G., Marker, L., Marnewick, K., Klein, R. and Williams, S. (2007). Regional assessment of the status, distribution and conservation needs of cheetahs in southern Africa. Cat News, 3, 44-46.

Ray, R.R. (2011). Ecology and population status and the impact of trophy hunting of the leopard *Panthera pardus* (Linnaeus, 1758) in the Luambe National Park and surrounding Game Management Areas in Zambia. [Doctoral dissertation, Rheinische Friedrich-Wilhelms-Universität].

Roberts, P.J., Druce, D.J., Mgqatsa, N. and Parker, D.M. (2024). Using photo by-catch data to reliably estimate spotted hyaena densities over time. Mammalia, 88(2), 85-92.

Rödel, H.G., Ibler, B., Ozogány, K. and Kerekes, V. (2023). Age-specific effects of density and weather on body condition and birth rates in a large herbivore, the Przewalski's horse. Oecologia, 203(3), 435-451.

Roth, T.L., Bateman, H.L., Kroll, J.L., Steinetz, B.G. and Reinhart, P.R. (2004). Endocrine and ultrasonographic characterization of a successful pregnancy in a Sumatran rhinoceros (*Dicerorhinus sumatrensis*) supplemented with a synthetic progestin. Zoo Biology, 23(3), 219-238.

Roth, T.L., Reinhart, P.R., Romo, J.S., Candra, D., Suhaery, A. and Stoops, M.A. (2013). Sexual maturation in the Sumatran rhinoceros (*Dicerorhinus sumatrensis*). Zoo Biology, 32(5), 549-555.

Santini, L., Isaac, N.J. and Ficetola, G.F. (2018). TetraDENSITY: A database of population density estimates in terrestrial vertebrates. Global Ecology and Biogeography, 27(7), 787-791.

Sargent, R. (2016). Investigating the effects of grassland management techniques on vegetation and wildlife at Lewa Wildlife Conservancy, Kenya. [MSc dissertation, University of Southampton].

Schaller, G.B. (1998). Wildlife of the Tibetan steppe. University of Chicago Press.

Shah, N. and Qureshi, Q. (2007). Social organization and determinants of spatial distribution of Khur (*Equus hemionus khur*). Erforschung biologischer Ressourcen der Mongolei, 10, 189-200.

Sharma, B.D., Clevers, J., De Graaf, R. and Chapagain, N.R. (2004). Mapping *Equus kiang* (Tibetan wild ass) habitat in Surkhang, upper mustang, Nepal. Mountain Research and Development 24, 149-156.

Shrestha, B.B., Lamichhane, B.R. and Amin, R. (2025). Population size, density, and ranging behaviour of a key leopard population in Nepal. Journal for Nature Conservation, 86, 126920.

Shrotriya, S., Reshamwala, H.S., Lyngdoh, S., Jhala, Y.V. and Habib, B. (2022). Feeding patterns of three widespread carnivores—the wolf, snow leopard, and red fox—in the Trans-Himalayan landscape of India. Frontiers in Ecology and Evolution, 10, 815996.

Sinclair, A.R.E., Mduma, S.A. and Arcese, P. (2000). What determines phenology and synchrony of ungulate breeding in Serengeti? Ecology, 81(8), 2100-2111.

Smielowski, J.M. and Raval, P.P. (1988). The Indian wild ass—wild and captive populations. Oryx, 22(2), 85-88.

Smuts, G.L. (1976). Reproduction in the zebra mare *Equus burchelli antiquorum* from the Kruger National Park. Koedoe, 19(1), 89-132.

Smyth, L.K., Rogan, M.S., Balme, G.A. and O'riain, M.J. (2025). Counting spots: Leopard density along a gradient of conservation rigor. Conservation Science and Practice, 7(12), e70197.

Somaweera, R., Brien, M. and Shine, R. (2013). The role of predation in shaping crocodilian natural history. Herpetological Monographs, 27(1), 23-51.

St-Louis, A. and Côté, S.D. (2009). Equus kiang (Perissodactyla: Equidae). Mammalian Species, 835, 1-11.

Stander, P.E. (1991). Demography of lions in the Etosha National Park, Namibia. Madoqua, 18(1), 1-9.

Stein, A.B., Fuller, T.K., DeStefano, S. and Marker, L.L. (2011). Leopard population and home range estimates in north-central Namibia. African Journal of Ecology, 49(3), 383-387.

Subedi, N., Lamichhane, B.R., Amin, R., Jnawali, S.R. and Jhala, Y.V. (2017). Demography and viability of the largest population of greater one-horned rhinoceros in Nepal. Global Ecology and Conservation, 12, 241-252.

Sundaresan, S.R., Fischhoff, I.R., Dushoff, J. and Rubenstein, D.I. (2007). Network metrics reveal differences in social organization between two fission–fusion species, Grevy's zebra and onager. Oecologia, 151(1), 140-149.

Talukdar, B.K. (2002). Tiger predation of rhino calves at Kaziranga National Park, Assam. Tiger Paper, 29(4), 19-21.

Thel, L., Bonenfant, C. and Chamaillé-Jammes, S. (2025). Good moms: dependent young and their mothers cope better than others with longer dry season in plains zebras. Oecologia, 207(3), 45.

Trinkel, M. (2009). A keystone predator at risk? Density and distribution of the spotted hyena (*Crocuta crocuta*) in the Etosha National Park, Namibia. Canadian Journal of Zoology, 87(10), 941-947.

Trinkel, M. (2013). Climate variability, human wildlife conflict and population dynamics of lions *Panthera leo*. Naturwissenschaften, 100(4), 345-353.

Trinkel, M., Ferguson, N., Reid, A., Reid, C., Somers, M., Turelli, L. et al. (2008). Translocating lions into an inbred lion population in the Hluhluwe-iMfolozi Park, South Africa. Animal Conservation, 11(2), 138-143.

Truter A. (2021). The reproductive performance, demography and spatial ecology of an extralimital white rhinoceros population. [MSc dissertation, Rhodes University].

Tumursukh, L., Suryawanshi, K.R., Mishra, C., McCarthy, T.M. and Boldgiv, B. (2016). Status of the mountain ungulate prey of the Endangered snow leopard *Panthera uncia* in the Tost Local Protected Area, South Gobi, Mongolia. Oryx, 50(2), 214-219.

Tupper, E. (2011). Foal survival and resource ecology of lactating Grevy's Zebra (*Equus grevyi*) Females. [Msc dissertation, University of Columbia].

Utete, B. (2021). A review of the conservation status of the Nile crocodile (*Crocodylus niloticus* Laurenti, 1768) in aquatic systems of Zimbabwe. Global Ecology and Conservation, 29, e01743.

van Der Meer, E. (2011). Is the grass greener on the other side?: testing the ecological trap hypothesis for African wild dogs (*Lycaon pictus*) in and around Hwange National Park. [Doctoral dissertation, Université Claude Bernard-Lyon I].

van der Meer, E. (2018). Carnivore conservation under land use change: the status of Zimbabwe's cheetah population after land reform. Biodiversity and Conservation, 27(3), 647-663.

Van Duyne, C., Ras, E., De Vos, A.E., De Boer, W.F., Henkens, R.J. and Usukhjargal, D. (2009). Wolf predation among reintroduced przewalski horses in Hustai National Park, Mongolia. The Journal of Wildlife Management, 73(6), 836-843.

Van Dyk, G. and Slotow, R. (2003). The effects of fences and lions on the ecology of African wild dogs reintroduced to Pilanesberg National Park, South Africa. African Zoology, 38(1), 79-94.

Van Strien, N.J. (1985). The Sumatran rhinoceros in the Gunung Leuser National Park, Sumatra, Indonesia: its distribution, ecology and conservation. [Doctoral dissertation, Landbouwhogeschool te Wageningen].

Wallace, K.M., Leslie, A.J., Coulson, T. and Wallace, A.S. (2013). Population size and structure of the Nile crocodile *Crocodylus niloticus* in the lower Zambezi valley. Oryx, 47(3), 457-465.

Webber, Q.M. and McGuire, L.P. (2022). Heterothermy, body size, and locomotion as ecological predictors of migration in mammals. Mammal Review, 52(1), 82-95.

Weise, F.J., Vijay, V., Jacobson, A.P., Schoonover, R.F., Groom, R.J., Horgan, J. *et al.* (2017). The distribution and numbers of cheetah (*Acinonyx jubatus*) in southern Africa. PeerJ, 5, e4096.

Whittington-Jones, B.M., Parker, D.M., Bernard, R.T. and Davies-Mostert, H.T. (2014). Habitat selection by transient African wild dogs (*Lycaon pictus*) in northern KwaZulu-Natal, South Africa: implications for range expansion. South African Journal of Wildlife Research, 44(2), 135-147.

Whittington, J. and Sawaya, M.A. (2015). A comparison of grizzly bear demographic parameters estimated from non-spatial and spatial open population capture-recapture models. PloS one, 10(7), e0134446.

Williams, S.D. (1998). Grevy's zebra: Ecology in a heterogenous environment. [Doctoral dissertation, University of London].

Williams, S.T. (2011). The impact of land reform in Zimbabwe on the conservation of cheetahs and other large carnivores. [Doctoral dissertation, Durham University].

Zhang, W., Fan, Z., Han, E., Hou, R., Zhang, L., Galaverni, M. *et al*. (2014). Hypoxia adaptations in the grey wolf (*Canis lupus chanco*) from Qinghai-Tibet Plateau. PLoS genetics, 10(7), e1004466.

Zhang, Y., Tariq, A., Hughes, A.C., Hong, D., Wei, F., Sun, H. *et al*. (2023). Challenges and solutions to biodiversity conservation in arid lands. Science of the Total Environment, 857, 159695.
